# Requirement for Fc effector function is overcome by binding potency for broadly reactive anti-alphavirus antibodies

**DOI:** 10.1101/2024.11.03.619087

**Authors:** Victoria Callahan, Matthew S. Sutton, Christina L. Gardner, Doreswamy Kenchegowda, Megan M. Dunagan, Mrunal Gosavi, Courtney Green, Tammy Y. Chen, Jessica Prado-Smith, Daniel Long, Jodi L. Vogel, Thomas M. Kristie, Chad S. Clancy, Crystal W. Burke, Mario Roederer, Julie M. Fox

## Abstract

Alphaviruses are emerging public health threats. Broadly reactive anti-alphavirus monoclonal antibodies (mAbs) have been shown to be protective in mouse models of infection. However, the mechanism of Fc-dependent or Fc-independent heterologous protection remains ill-defined *in vivo*. Here, we used two vaccine-elicited, broadly reactive, anti-alphavirus mAbs, SKT05 and SKT20, to establish correlates of mAb-mediated protection during Venezuelan equine encephalitis virus (VEEV) challenge. SKT20 required Fc effector functions to prevent lethality. In contrast, SKT05-mediated survival was independent of Fc effector functions, which is likely linked to early viral control through potent egress inhibition. However, control of virus replication and spread with SKT05 was Fc-dependent; these findings extended to additional *in vivo* models with alternative VEEV subtypes and chikungunya virus. During therapeutic delivery of SKT05, Fc effector functions were only required at 3 days post-infection. The necessity of Fc effector functions for SKT20 was related to mAb binding avidity rather than epitope and could be overcome by increasing the dose of SKT20 relative to the functional avidity of SKT05. Collectively, this study identified antibody avidity as a correlate for *in vivo* efficacy and associated Fc-dependent mechanisms that can be leveraged for therapeutic development of monoclonal antibodies against alphaviruses.

**One sentence summary:** Functional avidity of broadly reactive anti-alphavirus antibodies dictates requirement for Fc-mediated protection.

## INTRODUCTION

Alphaviruses, which belong to the family *Togaviridae*, are emerging and re-emerging mosquito-transmitted viruses of global concern. Alphaviruses are grouped based on geographic origin or symptomatic presentation into the New World (NW), or encephalitic, and Old World (OW), or arthritogenic, alphaviruses. The OW alphaviruses, including chikungunya virus (CHIKV), can cause arthritis and acute to chronic musculoskeletal disease. While CHIKV is globally distributed, the other OW alphaviruses are more geographically isolated. The NW alphaviruses, including Venezuelan equine encephalitis virus (VEEV), eastern equine encephalitis virus (EEEV), and western equine encephalitis virus (WEEV), circulate in North, Central, and South America, and infection can result in a range of symptoms from acute febrile illness to severe and potentially lethal encephalitis.

Alphaviruses circulate in enzootic and epizootic cycles using mosquitos and a variety of animal reservoirs. While transmission to humans is predominantly through mosquitoes, the encephalitic alphaviruses pose a potential biothreat due to ease of aerosol infection. Accordingly, VEEV, EEEV, WEEV are classified as Category B priority pathogens by the National Institutes of Health. Compounded by vector range expansion, geographical overlap of endemic regions, global movement of humans, and the absence of FDA-approved therapeutics, there is a need to develop broadly effective vaccines and therapeutics that can provide pan-alphavirus protection.

Alphaviruses have a positive-sense RNA genome of approximately 11.5 kb. The genome is comprised of four nonstructural (nsP1-4) and six structural proteins (capsid, E3, E2, 6K, TF, and E1) that are encoded by two open reading frames (*1, 2*). The mature virion is comprised of a nucleocapsid, surrounded by an envelope containing heterodimers of glycoproteins, E2 and E1, arranged as trimeric spikes. E1 and E2 are critical for receptor binding and attachment; and E1 has a hydrophobic fusion loop that aids in viral membrane fusion within the endosome (*1, 2*). E1 and E2 are also the primary antigenic targets of neutralizing antibodies (*2, 3*). Protective monoclonal antibodies (mAbs) have been identified against multiple alphaviruses that target E1 and E2 and can inhibit various steps in the viral life cycle, such as attachment, entry, fusion, and egress (*3–7*). Antibodies clear infected cells and opsonized virions, as well as modulate the immune response through Fc mediated effector functions (*2, 8–10*). However, few studies evaluated the requirement of neutralization and/or Fc effector functions for optimal pan-alphavirus protection.

We previously identified two broadly reactive anti-alphavirus mAbs, SKT05 and SKT20, isolated from cynomolgus macaques vaccinated with a mix of VEEV, WEEV, and EEEV virus like-particles (VLPs) (*11*). SKT05 and SKT20 bound to NW and OW alphaviruses but had differential neutralization of Env-pseudotyped lentivirus reporter viruses expressing the glycoproteins of VEEV, EEEV, or WEEV (*11*). Both SKT05 and SKT20 protected mice from a lethal VEEV (strain TC-83) challenge. However, only SKT05 reduced viral loads in the brain (*11*). While both antibodies engage residues near or at the highly conserved E1 fusion loop, SKT05 and SKT20 bound to distinct, non-competing epitopes and with different angles of approach (*11*). Still, the mechanism(s) of protection afforded by SKT05 and SKT20 and the antibody characteristics (*e.g.,* avidity or epitope) that correlated with viral control *in vivo* has not been defined.

Here, we used a lethal mouse model of VEEV-induced encephalitis to elucidate the mechanism of protection for SKT05 and SKT20 through evaluation of viral kinetics, the inflammatory response, and histopathology using mAb variants with minimal binding to Fc gamma receptors (FcγRs) and the complement component, C1q. We find that Fc effector functions were not required for SKT05-mediated survival, but necessary for late control of viral burden. Similar phenotypes were observed with additional VEEV subtypes and CHIKV. SKT05 controlled early viral replication through egress inhibition and Fc effector functions were dispensable for SKT05 therapeutic protection up to two days post-infection (dpi) with VEEV. In contrast, for SKT20, Fc engagement was necessary to reduce a pro-inflammatory response and provide protection. Potency of Env-pseudovirus neutralization and binding to infected cells related to the requirement of Fc-mediated protection. Indeed, the Fc-dependency of SKT20 *in vivo* could be overcome by administering an equivalent dose based on SKT05 potency *in vitro*. Overall, this study lends novel insight to predictors of *in vivo* efficacy and mechanism of broadly reactive, anti-alphavirus antibodies.

## RESULTS

### Broadly reactive anti-alphavirus mAbs protect against lethal VEEV challenge

Prophylactic administration of SKT05 and SKT20 prevented mortality in C3H/HeN mice intranasally inoculated with the BSL-2 vaccine strain of VEEV, TC-83 (*11*). Numerous studies have demonstrated that the C3H/HeN TC-83 model results in encephalitis, similar to models with virulent strains of VEEV (*12, 13*). We sought to verify the protective effects of SKT05 in a fully virulent model of VEEV. BALB/c mice were administered SKT05, an anti-E1 VEEV-specific mAb (SKV09), a positive control (1A3B-7), or a control rhesus mAb (ITS103.01) one hour post aerosol challenge with a lethal dose of VEEV Trinidad Donkey (TrD; subtype IAB). SKT05-treated mice survived and showed minimal weight-loss and clinical disease similar to mice treated with SKV09 or positive control (**Fig. 1A-C**). With consistent findings between the models (**Fig. 1A and D**), we proceeded with the C3H/HeN model of VEEV TC-83 induced encephalitis via intranasal inoculation.

**Figure 1.**
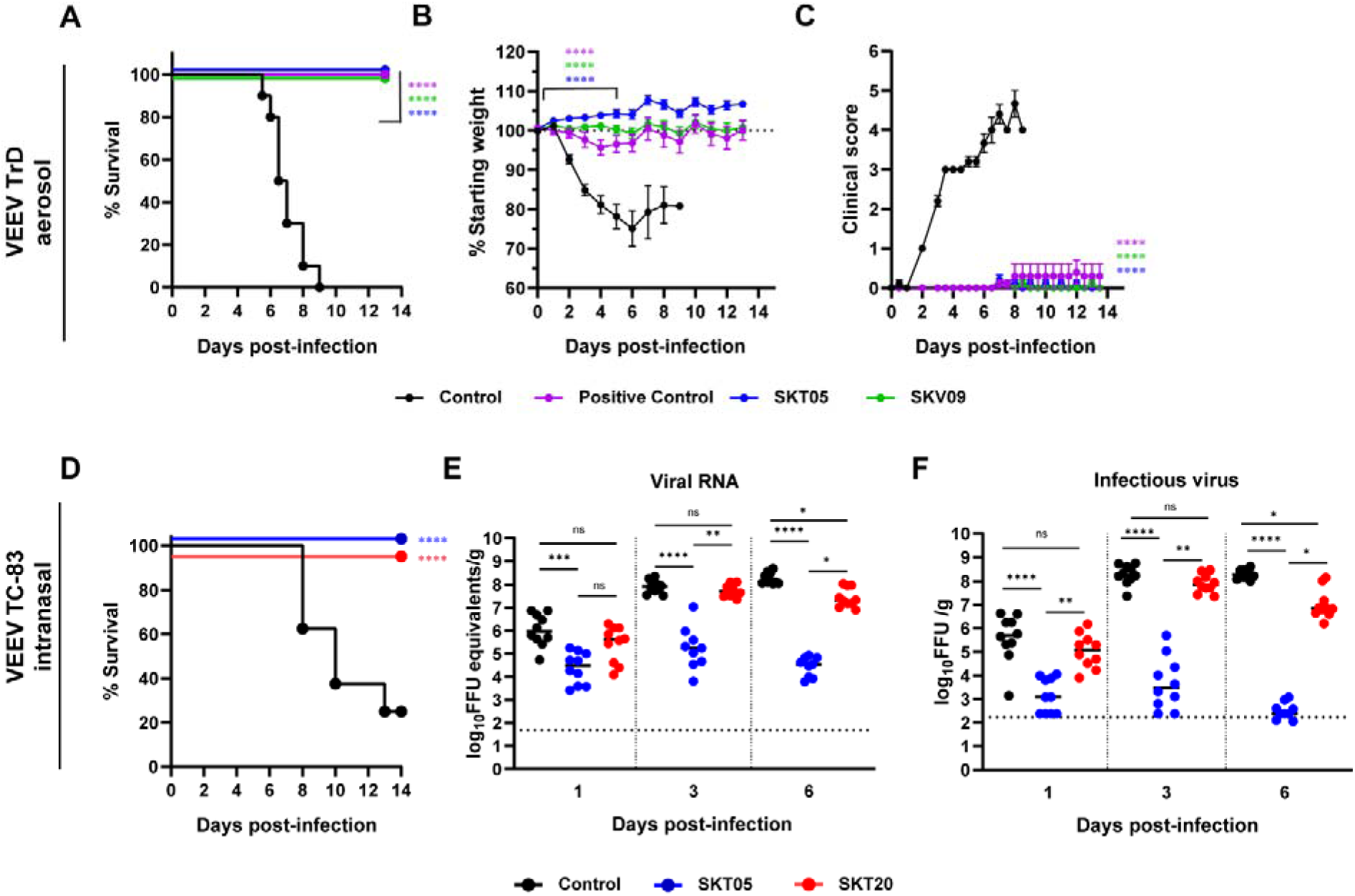
Broadly reactive anti-alphavirus mAbs protect against lethal VEEV challenge. (**A-C**) BALB/C were administered 200 µg of SKT05, SKV09, a positive control antibody (1A3B7), or control antibody at 1 h post-aerosol challenge with 10^3^ PFU of VEEV strain Trinidad Donkey (TrD). Mice were monitored for 14 days for survival (A), weight-loss (mean ±SEM) (B), and clinical score (mean ±SEM) (C) (n = 10 mice/group; 2 independent experiments). (**D-F**) C3H/HeN mice received 200 µg of indicated mAb 1 day prior to intranasal challenge with 10^7^ FFU of VEEV strain TC-83 (n = 10/group; 2 independent experiments). (D) Survival was followed for 14 dpi. Brains were harvested from a separate group of mice at 1, 3, and 6 dpi and viral RNA (E) or infectious virus (F) was determined by RT-qPCR or FFA, respectively. Bars represent the median. Statistical significance was determined by a Log-rank test (A, D), one-way ANOVA with a Dunnett’s post-test of area under the curve (AUC) analysis from 0-5 dpi comparing each treatment to the control group (B-C), and Kruskal-Wallis with a Dunn’s post-test comparing all groups (E-F). The dotted line indicates starting body weight (B) or the limit of detection (LOD; E-F).

To determine the early impact of SKT05 and SKT20 on VEEV infection, mAbs were administered one day before TC-83 infection and viral loads were measured in the brain, serum, and spleen at 1, 3, and 6 dpi. SKT05 and SKT20 reduced viral RNA and infectious virus in the brain at 1 dpi (**Fig. 1E-F**), which correlated with reduced viral RNA in the serum and spleen (**Fig. S1A-B**). However, only SKT05 treatment controlled viral burden across all assessed time-points. The viral load in the brain of SKT20-treated mice was comparable to control-treated mice at 3 dpi but began to decrease by 6 dpi (**Fig. 1E-F**).

While both mAbs reduced virus early, SKT05 was superior at controlling virus in the brain. This could not be explained by differences in antibody bioavailability as there were no differences in the quantity of SKT05 and SKT20 in the brain or serum of mice at 5 dpi (**Fig. S1C**). Our previous study demonstrated that VEEV Env-pseudotyped viruses are differentially neutralized by SKT05 (IC_50_: 0.01 µg/mL) and SKT20 (IC_50_: 0.19 µg/mL) (**Fig. S1D**). Another explanation for the differential protection *in vivo* may be increased neutralization of TC-83 by SKT05. However, SKT05 and SKT20 failed to or poorly neutralized TC-83, respectively, while neutralization was observed with an anti-E1 VEEV-specific mAb, SKV09 (IC_50_: 0.15 µg/mL) (**Fig. S1E**). These results suggest that SKT05 and SKT20 use distinct mechanisms to mediate VEEV protection in mice.

### SKT05 and SKT20 reduce pro-inflammatory cytokines and chemokines during VEEV infection

Poor clinical outcome and lethality are not directly correlated to viral load in the brains of VEEV-infected mice. Indeed, neuroinvasion precedes blood brain barrier (BBB) disruption and the development of encephalitis (*14*). Additionally, BBB disruption coincides with elevated numbers of infiltrating immune cells, loss of tight-junctions, and increased pro-inflammatory cytokine and chemokine expression (*14–18*). Intracellular adhesion molecule -1 (ICAM-1), which promotes leukocyte adhesion and transmigration, is increased, in conjunction with, the altered localization of proteins that regulate permeability and integrity of the brain microvascular endothelium (*19–21*). Given the differences in viral load in the brain, we analyzed transcriptional signatures that correlate with BBB disruption by RT-qPCR from brains of mAb-treated mice at 5 dpi (**Fig. 2A**). SKT05-treated mice had reduced expression of ICAM-1, C-X-C motif chemokine ligand (CXCL) 9 (CXCL9), and CXCL10 gene-transcripts compared to both control and SKT20-treated mice (**Fig. 2A**). Notably, SKT20-treated mice upregulated ICAM-1, CXCL9, and CXCL10 and the levels were not statistically different from control mice. There was no difference in matrix metalloproteinase 9 (MMP-9) gene-expression between the treatment groups at this time-point (**Fig. 2A**).

**Figure 2.**
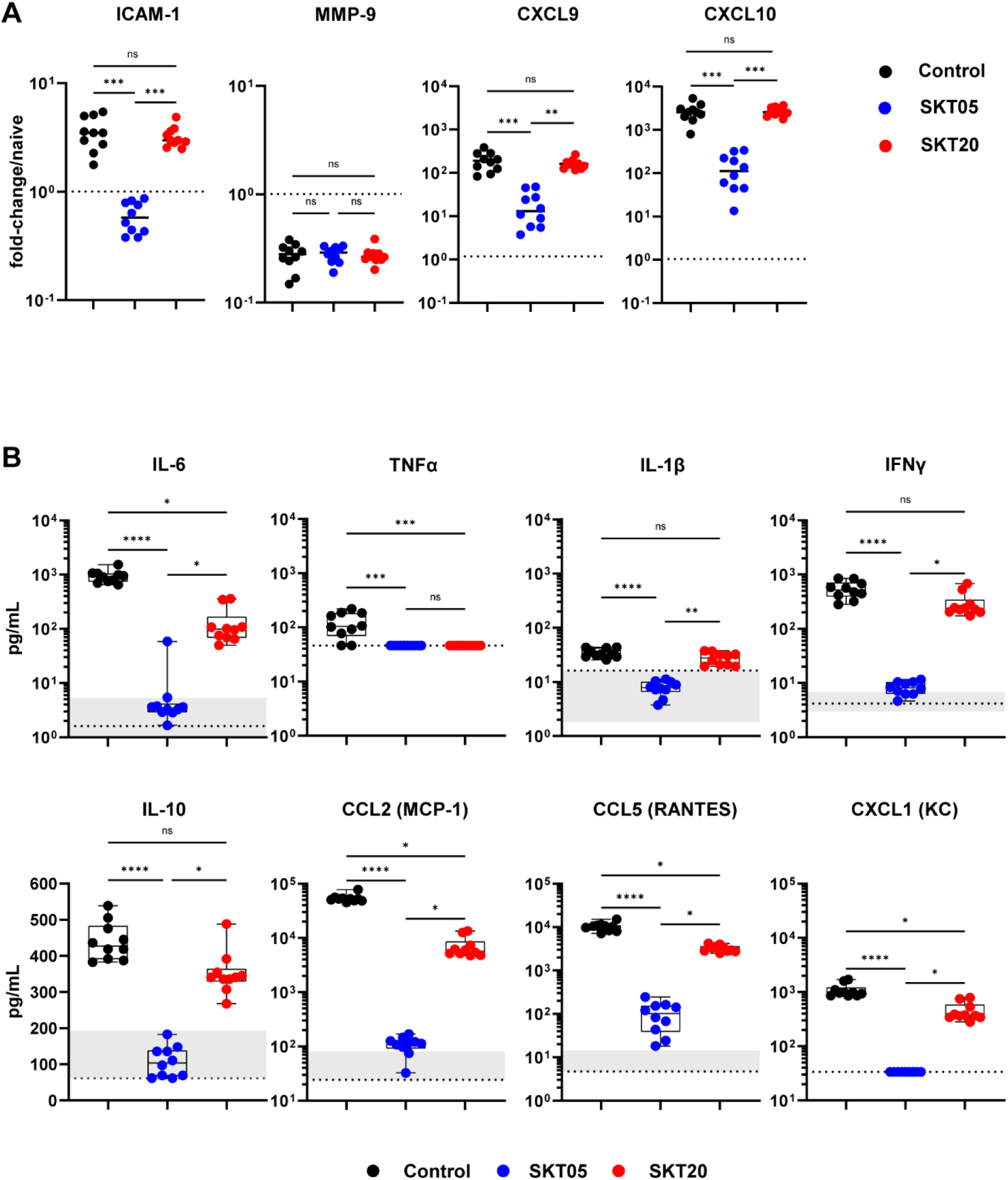
SKT05 and SKT20 reduce pro-inflammatory cytokines and chemokines during VEEV infection. C3H/HeN were administered 200 µg of indicated mAb at 1 day before infection with TC-83. (**A**) Brains were harvested at 5 dpi for gene-expression analysis by RT-qPCR (n = 10/group; 2 independent experiments). Samples were normalized to *Gapdh* and then compared to naïve tissue. Fold-change was calculated using 2^-ΔΔCt^. The dotted line represents the average fold-change in naïve mouse controls (n = 5). Bars represent the median. (**B**) Brains were harvested at 6 dpi and the concentration of cytokines and chemokines were determined using a Bio-plex assay. The box and whisker indicate the min to max (n = 10/group; 2 independent experiments). The min and max concentration of the analyte detected in the naïve brain homogenate is represented by the shaded regions within the graphs. The dotted line represents the LOD of the assay. If naïve samples were at the LOD, only a dotted line is shown. (A-B) Statistical significance was determined by Kruskal-Wallis with a Dunn’s post-test comparing all groups.

We next evaluated the level of pro-inflammatory cytokines and chemokines which contribute to inflammation and immune cell recruitment. Cytokines, like TNF-α, have been associated with VEEV-induced BBB permeability and neurodegeneration (*22*). Other cytokines and chemokines such as IL-6, IFNγ, IL-1β, C-C motif ligand (CCL) 2 (CCL2), and CCL5 have positive correlation to VEEV-induced pathological findings (*23*). Mice were treated with SKT05, SKT20, or a control mAb one day before TC-83 infection. At 6 dpi, brains were harvested, and pro-inflammatory cytokines and chemokines were analyzed by a Bio-Plex assay (**Fig. 2B, Table S1**). Naïve mice were included in the analysis to determine baseline levels. Overall, SKT05 treatment dramatically reduced proinflammatory cytokines and chemokines compared to control and SKT20-treated mice (**Fig. 2B**). Notably, IL-6, TNF-α, IL-1β, and IFNγ were reduced to levels observed in naïve mice. Other cytokines like IL-10, as well as chemokines CCL2, CCL5, and CXCL1, were also significantly reduced with SKT05 treatment. SKT20 reduced IL-6, TNFα, CCL2, CCL5, and CXCL1 compared to control treated mice, albeit to a lesser degree than SKT05 (**Fig. 2B**). These results suggest that SKT05 treatment dampens the inflammatory environment, presumably due to limited virus within the brain, while a heightened inflammatory environment was evident with SKT20 treatment suggesting robust immune cell recruitment.

### Fc effector functions are required for SKT20 protection but dispensable for SKT05-mediated survival during VEEV challenge

Antibodies can reduce alphavirus infection and disease *in vivo* by blocking entry, fusion, or egress of viral particles and through interactions of the Fc region with Fc receptors (*4–9, 24, 25*). Since SKT05 and SKT20 lack robust TC-83 neutralization *in vitro,* we assessed the necessity of Fc effector functions for *in vivo* protection. We generated LALA-PG variants of SKT05 and SKT20, which abrogate binding to FcγRs and C1q (*26*). First, we confirmed equivalent neutralization of the wild-type and LALA-PG antibodies against VEEV pseudovirus in cell culture (**Fig. 3A**). Then, we verified that the LALA-PG mAbs fail to appreciably bind to the human high affinity FcγR, FcγRI, by ELISA (**Fig. 3B**). Next, mice were administered wild-type or LALA-PG variants of SKT05 or SKT20 one day prior to TC-83 infection and were followed for weight-loss (**Fig. 3C-D**) and survival (**Fig. 3E-F**). Notably, SKT05-mediated survival was not dependent on Fc effector functions (**Fig. 3E**), although SKT05 LALA-PG-treated mice lost about 10% of starting weight (**Fig. 3C**). In contrast, SKT20-mediated protection was dependent on Fc-engagement as SKT20 LALA-PG-treated mice lost weight comparable to the control mice (**Fig. 3D**) and only 25% of the mice survived (**Fig. 3F**). These results indicate that SKT20 protects against VEEV lethality through Fc effector functions while SKT05 primarily protects through an Fc-independent mechanism.

**Figure 3.**
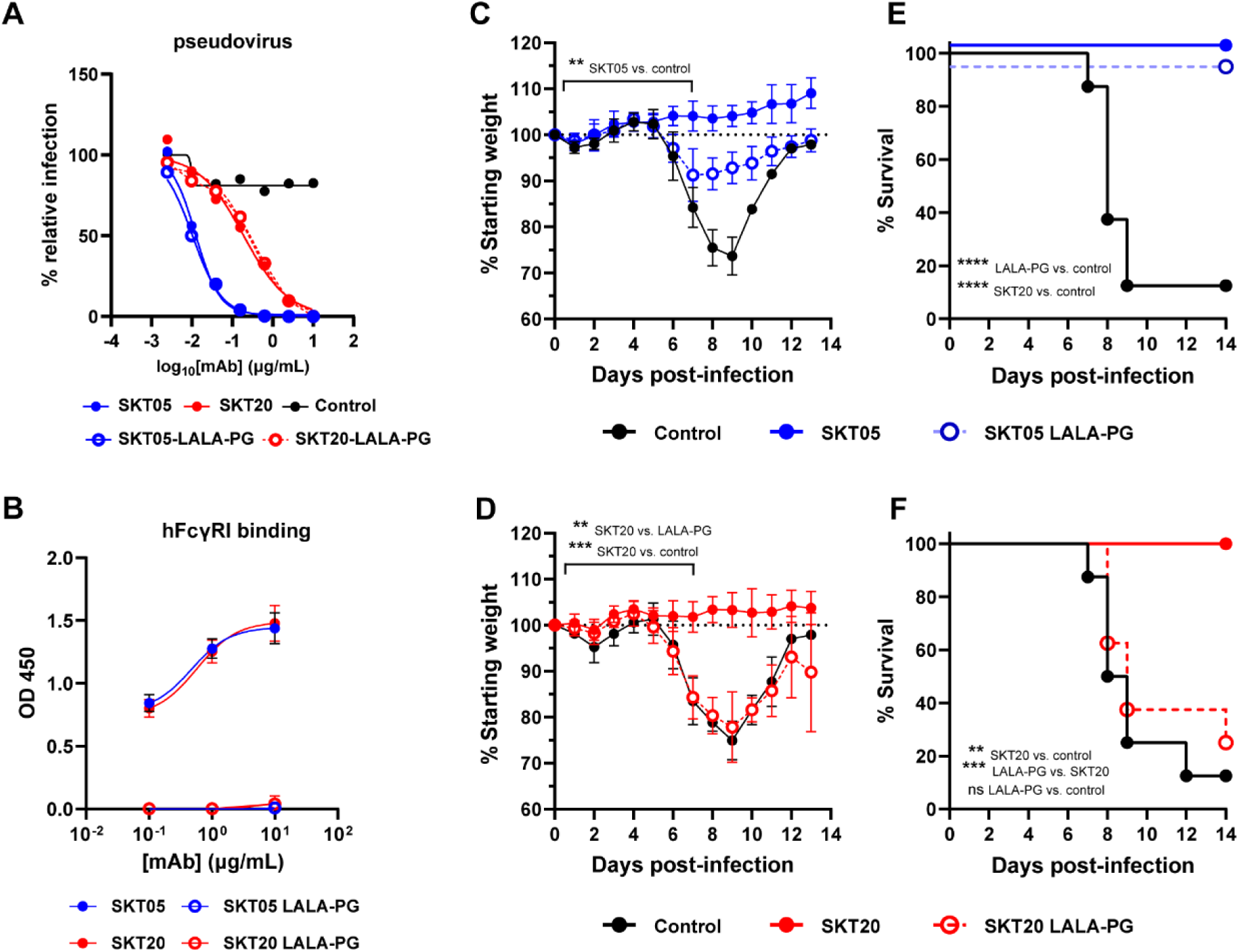
Fc effector functions are required for SKT20 protection but dispensable for SKT05-mediated survival during VEEV challenge. (**A**) Neutralization curve for wild-type and LALA-PG variants against VEEV Env-pseudotyped virus. Data is representative of one independent experiment, conducted in triplicate. (**B**) Binding of indicated mAbs to purified human (h) FcγRI was assessed by ELISA. The data is represented as the mean ± SD of two independent experiments performed in duplicate. (**C-F**) Indicated mAbs (200 µg) were administered to C3H/HeN mice 1 day prior to inoculation with TC-83. Mice were followed for weight loss (mean ±SD) (C, D) and survival (E, F) for 13-14 days (n = 8/group; 2 independent experiments). Statistical significance was determined by a one-way ANOVA with a Tukey’s post-test of AUC analysis from 0-7 dpi comparing all groups (C and D) or a Log-rank test (E and F).

### SKT20 alters the pro-inflammatory response and immune cell infiltrates in the brain through Fc effector functions

To determine whether SKT20 reduced viral burden and inflammation through Fc-engagement, mice were administered SKT20, SKT20 LALA-PG, or a control antibody, infected with TC-83, then viral RNA and pro-inflammatory cytokines and chemokines were measured in the brain at 6 dpi. As previously observed, SKT20 reduced viral RNA and levels of inflammatory cytokine and chemokines compared to the control treated mice (**Fig. 4A-B**). However, mice treated with SKT20 LALA-PG failed to clear viral RNA and had levels of pro-inflammatory cytokines and chemokines similar to control treatment (**Fig. 4A-B**).

**Figure 4.**
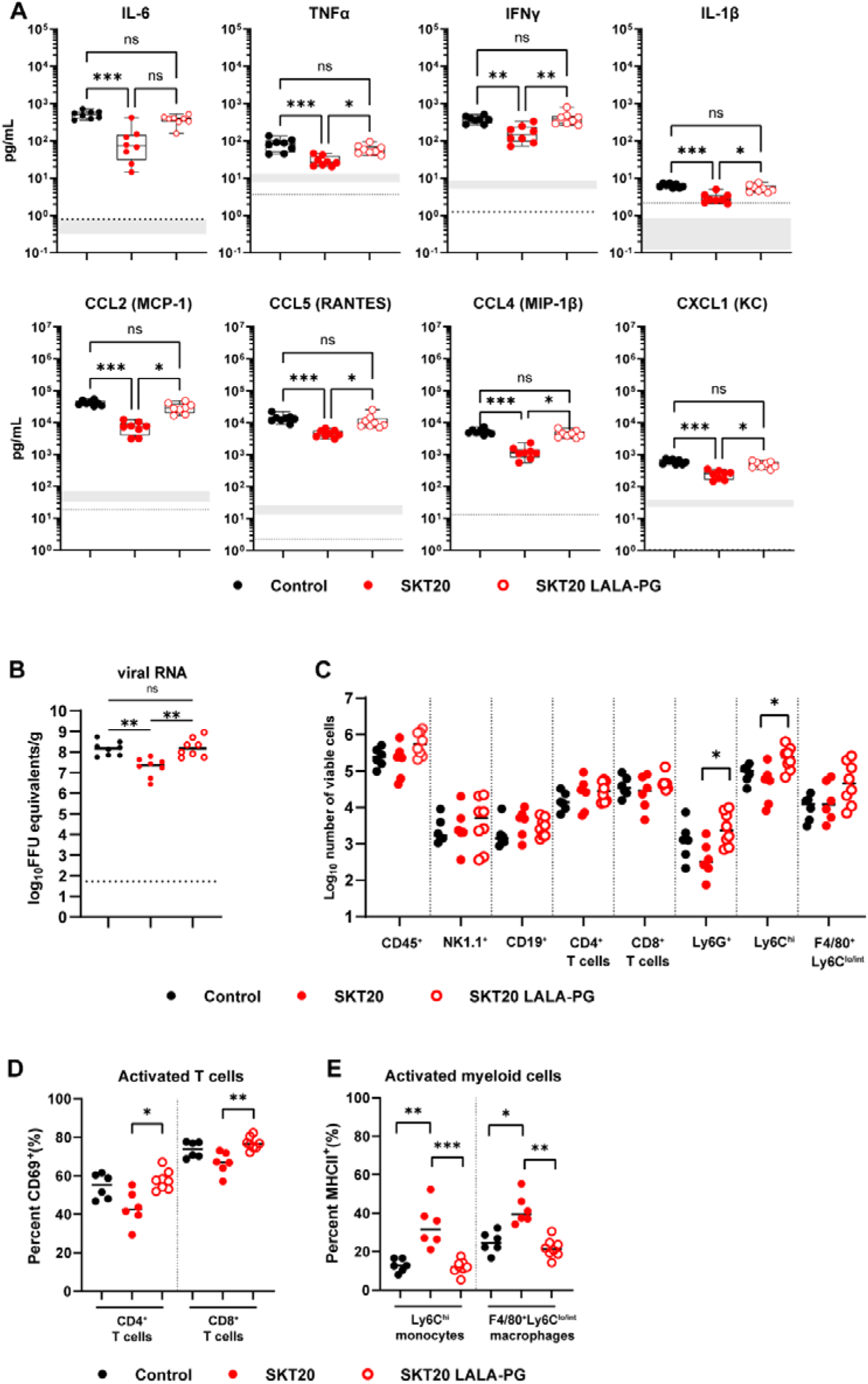
SKT20 alters the pro-inflammatory response and immune cell infiltrates in the brain through Fc effector functions. C3H/HeN mice were administered an isotype control, SKT20, or SKT20 LALA-PG (200 µg) at 1 day prior to infection with TC-83. (**A-B**) At 6 dpi, brains were harvested. (A) Chemokine and cytokine concentrations were determined using a Bio-plex assay. The box and whisker plot indicates the min to max (n=10/group, 2 independent experiments). The shaded bar represents the min and max concentration for naïve animals. The dotted line represents the LOD of the assay. If naïve samples were at the LOD, only a dotted line is shown. (B) Viral RNA in brain homogenate was determined by RT-qPCR (n = 6-8/group; 2 independent experiments). (**C-E**) At 6 dpi, brains were collected and digested. (C) Single cell suspensions were stained, and flow cytometry was performed to assess the total number of indicated cells. The percentage of activated (CD69^+^) CD4^+^ and CD8^+^ T cells (D) and activated (MHCII^+^) monocytes and macrophages (E) was determined by flow cytometry. (B-E) Bars represent the median. For all graphs, statistical significance was determined by Kruskal-Wallis with a Dunn’s post-test comparing all groups.

Previous work has shown an influx of monocytes, macrophages, and T cells into the brains of TC-83 infected mice between 3-7 dpi (*15, 20*). To determine if the modified chemokine response altered the infiltrating immune cell populations, we harvested brains from mAb-treated, TC-83 infected mice at 6 dpi and analyzed the immune cell subsets by flow cytometry (**Fig. S2A**). Analysis of NK cells (NK1.1^+^), B cells (CD19^+^), CD4^+^ T cells, CD8^+^ T cells, neutrophils (Ly6G^+^), macrophages (Ly6C^lo/int^ F4/80^+^), and monocytes (Ly6C^hi^) showed increased numbers of neutrophils and monocytes with SKT20 LALA-PG treatment compared to SKT20 (**Fig. 4C**). When the proportion of each population amongst the CD45^+^ cells was evaluated, SKT20 LALA-PG treated mice had a higher percentage of monocytes compared to SKT20 mice, while the percentage of CD4^+^ T cells was higher in SKT20-treated mice as compared to SKT20 LALA-PG treatment (**Fig. S2B**). However, the CD4^+^ and CD8^+^ T cells from SKT20-treated mice had reduced expression of the activation marker, CD69 (**Fig. 4D**). Notwithstanding, the total number of activated CD4^+^ and CD8^+^ T cells was similar between the groups (**Fig. S2C**). Myeloid cell populations were evaluated for MHCII expression as a marker of activation. Mice that received SKT20 had an increased percentage of activated monocytes and macrophages as compared to SKT20 LALA-PG or control mice although the total number of activated cells was similar across the groups (**Fig. 4E & S2D**). These results suggest that SKT20 may mediate clearance of viral RNA through Fc-FcγR interaction with monocytes and macrophages.

### SKT05 limits neuroinvasion and spread into caudal regions of the brain through VEEV egress inhibition

To determine if early viral control and reduced dissemination into the brain was dependent on Fc effector functions, we administered wild-type or a LALA-PG variant of SKT05 to mice one day before TC-83 challenge and harvested brains at 1 and 6 dpi. Both SKT05 and SKT05 LALA-PG reduced viral burden at 1 dpi (**Fig. 5A**). In contrast, reduction in viral RNA was highly dependent on Fc-engagement at 6 dpi (**Fig. 5A**).

**Figure 5.**
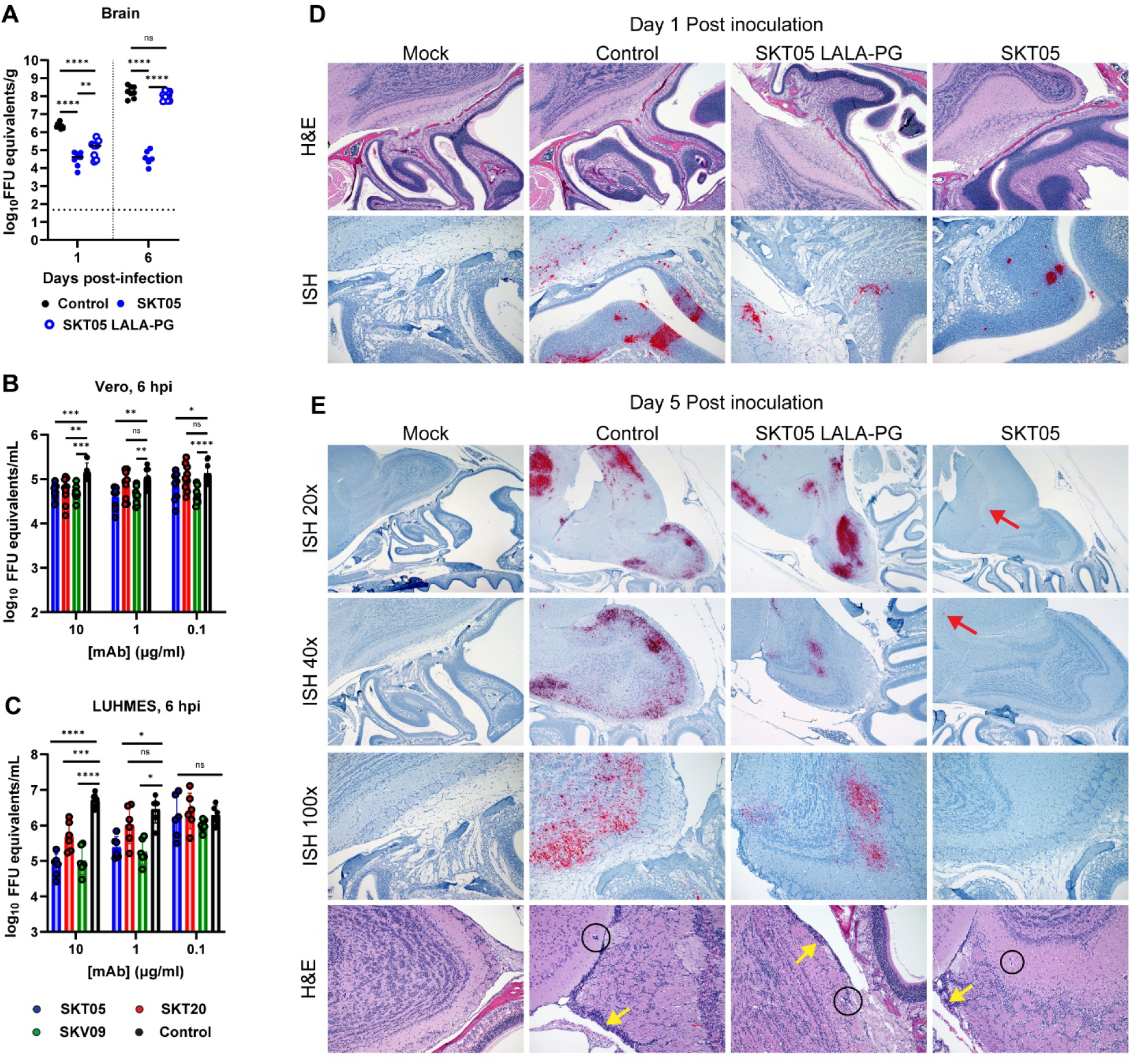
SKT05 limits neuroinvasion and spread into the brain through inhibition of viral egress. (**A**) C3H/HeN mice were treated with 200 µg of SKT05, SKT05-LALA-PG, or a control one day prior to infection with TC-83. At 1 and 6 dpi, viral loads were determined in the brains by RT-qPCR (n = 8/group; 2 independent experiments). The median is represented, and the dotted line indicates the LOD of the assay. Statistical significance was determined by one-way ANOVA with Holm-Sidak’s post-test. (**B-C**) Viral egress inhibition by indicated mAbs was evaluated in Vero cells (B) and LUHMES (C). Supernatants were collected at 6 hpi to quantify viral RNA by RT-qPCR. Data is representative of the mean ± SD of 2-3 independent experiments performed in triplicate. Statistical significance was determined by two-way ANOVA with Dunnett’s post-test comparing all groups to the isotype control. (**D-E**) C3H/HeN mice were pre-treated with 200 µg of SKT05, SKT05-LALA-PG, or a control antibody one day before infection with TC-83. At 1 and 5 dpi, skulls with brains intact were harvested, fixed, then decalcified before paraffin embedding and sectioning. Representative images of sagittal skull and brain sections stained for VEEV RNA by *in situ* hybridization (ISH) or with hematoxylin and eosin (H&E). The red arrows point out focal vRNA staining, the yellow arrows indicate meningitis, and the circles indicate perivascular cuffing. Data are representative images of two independent experiments (n = 6 - 8/group).

Our initial findings showed that TC-83 is not neutralized by SKT05 despite observed VEEV pseudovirus neutralization (**Fig. S1E**). Since early control of VEEV infection was Fc-independent, we evaluated SKT05 inhibition of different stages in the viral life cycle. Our standard focus reduction neutralization test (FRNT) allows for multiple rounds of VEEV replication which may permit VEEV to overcome early SKT05 entry inhibition. Entry inhibition was assessed using a single-cycle, viral entry inhibition assay. mAbs were pre-incubated with TC-83 then added to cells. Following extensive washing, cells were incubated in medium containing ammonium chloride to prevent *de novo* infection. VEEV antigen^+^ cells were determined by flow cytometry. SKV09, a VEEV specific mAb that binds E1 and neutralizes TC-83, was included as a positive control (**Figure S1E**) (*11*). In Vero cells, SKT05 (IC_50_: 0.6388 µg/mL) and SKT20 (IC_50_: 1.551 µg/mL) showed some level of entry inhibition compared to a control mAb (**Fig. S3A, *left***) while SKV09 provided nearly complete entry inhibition with only 3% relative infection observed (**Fig. S3A, *left***). Neuronal cells are one of the cellular targets of VEEV infection via both hematogenous and intranasal routes of infection (*27, 28*). We next used a neuronal cell line, Lund human mesencephalic cells (LUHMES), to determine if SKT05, SKT20, and SKV09 differentially inhibit entry in a natural target of VEEV infection. Surprisingly, there was minimal entry inhibition with all three antibodies, SKT05, SKT20, and SKV09 (**Fig. S3A, *right***).

We next assessed inhibition of viral egress. Vero cells or LUHMES were infected with TC-83, washed extensively, then serial dilutions of mAbs were added to the cells in medium containing ammonium chloride. Viral RNA level was determined in supernatants at 1 **(Fig. S3B-C)** and 6 hours post infection (hpi) (**Fig. 5B-C**). In Vero cells, SKT05 significantly inhibited egress in a dose-dependent manner at 6 hpi as compared to control treated cells (**Fig. 5B**). In contrast, SKT20 only reduced virus release at the highest concentration and to a lesser degree than SKT05 and SKV09. SKV09 showed a similar reduction in viral RNA levels across all concentrations (**Fig. 5B**). In the LUHMES, SKV09 and SKT05 showed a similar dose-dependent inhibition, while SKT20 only significantly reduced virus egress at the highest concentration (**Fig. 5C**). Importantly, SKT05 was superior to SKT20 in VEEV egress inhibition suggesting this may contribute to early VEEV protection *in vivo*.

Intranasal inoculation with VEEV results in initial infection of the olfactory neuroepithelium. The virus disseminates into the brain by anterograde transport along the axonal tracts through the cribriform plate and then seeds the olfactory bulb (*27*). CNS pathology in infected mice may include necrosis and apoptosis of neurons, endothelial cell injury, lymphocyte destruction, perivascular cuffing, and meningitis (*12, 14, 15, 17*). To determine if SKT05-mediated restriction of VEEV dissemination into the brain is Fc-dependent, we performed *in situ* hybridization (ISH) to probe for VEEV RNA and Hematoxylin and Eosin (H&E) staining on sequential midsagittal skull and brain sections from mice treated with SKT05, SKT05 LALA-PG, or a control antibody at 1 and 5 dpi. We chose 5 dpi for this study to observe a time point just prior to the onset of weight loss, so we could identify features that may promote disease. Mock-infected mice showed no significant histopathologic changes, and no detectable viral RNA labeling (**Fig. 5D-E**). At 1 dpi, control-treated mice had mild olfactory epithelial and olfactory nerve tract necrosis (**Fig. 5D and S3D**). Consistent with previous reports, control-treated mice had no observable viral RNA labeling in the respiratory epithelium, but abundant labeling in regions of olfactory epithelium, olfactory nerve tracts and within the olfactory bulb (**Fig. S3E**) (*29*). Similarly, SKT05 LALA-PG treated mice showed mild olfactory epithelial and peri-neural tract necrosis at 1 dpi. Viral RNA labeling was present in the olfactory epithelium, olfactory nerve tracts, and olfactory bulb, but at a more limited distribution compared to the control-treated mice. SKT05 treated mice displayed minimal necrosis of the olfactory epithelial and olfactory neural tracts. Viral labeling was observed at a low distribution in olfactory epithelium, olfactory neural tracts and in half of evaluated sections of olfactory bulb.

By 5 dpi, control-treated mice had moderate to severe neuroparenchymal necrosis with abundant necrotic neurons and multifocal infiltrates of degenerate leukocytes accompanied with multifocal, pleocellular meningitis and perivascular cuffing (**Fig. 5E and Fig. S3D**). Remarkably, only minimal changes were observed in the olfactory epithelium and olfactory neural tracts. *In-situ* hybridization revealed abundant viral RNA labeling throughout the olfactory bulb, cerebrum, and brainstem (**Fig. S3E**). SKT05 LALA-PG-treated mice showed moderate necrosis and meningitis in the olfactory bulb with perivascular cuffing. Abundant viral RNA was labeled throughout the olfactory bulb, cerebrum, and brainstem. In stark contrast, SKT05-treated mice exhibited minimal histopathologic changes in the brain. Similarly, viral RNA was rarely observed with limited, small foci visible in either the olfactory bulb or cerebrum. These results show that SKT05 Fc effector functions are necessary to prevent spread of virus into the brain while SKT05-mediated inhibition of viral egress aids in reducing the initial infection in the olfactory epithelium.

### Fc effector function necessity is dependent on functional avidity rather than epitope specificity

While SKT05 and SKT20 both bind to the E1 protein and are broadly reactive, they interact with distinct, non-competing epitopes and differ in their VEEV pseudovirus neutralization potency (**Fig. S1D**) (*11*). It is unclear whether the requirement of Fc effector functions for VEEV protection correlates with antibody epitope or avidity. To begin to evaluate avidity as a correlate of protection, we measured binding of SKT05 and SKT20 to the surface of live TC-83 infected Vero cells by flow cytometry. SKV09 was included as a positive control. SKT05 (EC_50_: 0.26 µg/mL) and SKV09 (EC_50_: 0.31 µg/mL) were the strongest binders while there was a roughly 10-fold difference in the binding potency of SKT20 (EC_50_: 2.8 µg/mL) (**Fig. 6A-C**). Notably, differences in binding to infected cells mirrored differences in VEEV pseudotyped virus neutralization where SKT05 and SKT20 IC_50_ values differed by about 15-fold (**Fig. S1D**).

**Figure 6.**
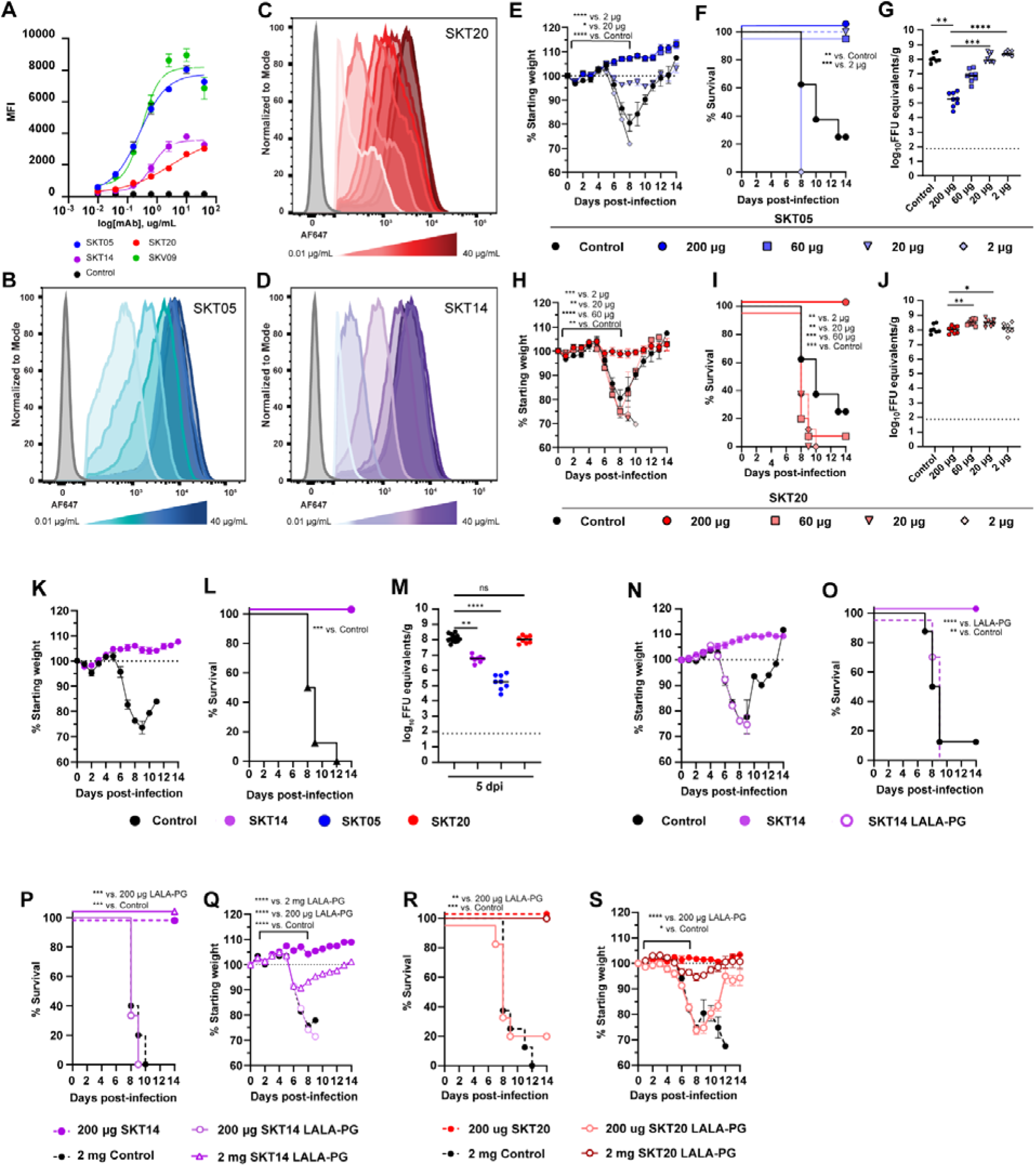
Fc effector function necessity is dependent on functional avidity rather than epitope specificity. (**A-D**) Binding of indicated mAbs to the surface of live Vero cells infected with TC-83 was determined by flow-cytometry. EC_50_ values were determined by nonlinear regression of log transformed data. Data shows the median of 2 independent experiments performed in duplicate. (B-D) Histograms show binding of dilutions of SKT05 (B), SKT20 (C), and SKT14 (D) to the surface of live, TC-83 infected Vero cells. (**E-J**) C3H/HeN mice were administered 200, 60, 20, or 2 µg of SKT05, SKT20, or 200 µg of a control, one day prior to infection with TC-83. Mice were followed for weight loss (E, H) and survival (F, I) for 14 days. (G, J) Viral loads were determined in the brains of mice at 5 dpi (n = 8/group; 2 independent experiments). Statistical significance was determined by a one-way ANOVA with a Dunnett’s post-test of AUC analysis from 0-8 dpi comparing each group to the 200 µg group (E and H), Log-rank test compared to 200 µg group (F and I), or Kruskal-Wallis with a Dunn’s post-test comparing each group to the 200 µg dose (G and J). (**K-O**) C3H/HeN mice were administered 200 µg of SKT14, SKT14 LALA-PG, or a control mAb one day prior to infection with TC-83. Mice were followed for weight loss (K and N) and survival (L and O). Additional mice were sacrificed at 5 dpi to determine viral loads (M) in brain tissues by RT-qPCR (data related to SKT05 and SKT20 is the same as the 200 µg doses of in G, J) (n = 8/group; 2 independent experiments). Statistical significance was determined by a Log-rank test compared to the SKT14 group (L and O) or Kruskal-Wallis with a Dunn’s post-test compared to the control group (M). (**P-S**) C3H/HeN mice were administered 200 µg or 2 mg of SKT14 LALA-PG or SKT20 LALA-PG, 200 µg of SKT14 or SKT20, or 2 mg of a control mAb one day prior to infection with TC-83. Mice were followed for survival (P, R) and weight-loss (Q, S) for 14 days (n = 8/group; 2 independent experiments). Statistical significance was determined by a Log-rank test (P and R) or a one-way ANOVA with a Dunnett’s post-test of AUC analysis from 0-7 dpi comparing each group to the 200 µg SKT14 or SKT20 group (Q and S). For viral loads, bars represent the median. Weight loss is shown as mean ± SEM.

To test this *in vivo*, we administered approximately half log decreasing doses (200 µg, 60 µg, 20 µg, or 2 µg) of SKT05 (**Fig. 6E-G**) or SKT20 (**Fig. 6H-J**) to mice prior to challenge with TC-83. Mice were followed for weight-loss and survival. Tissues were collected at 5 dpi from a separate set of mice for analysis of viral load. Mice provided 200, 60, or 20 µg of SKT05 had 100% survival (**Fig. 6F**) and only mice treated with the 20 µg dose lost minimal weight (**Fig. 6E**). In contrast, 2 µg of SKT05 failed to protect mice from a lethal challenge (**Fig. 6E-F**). There was a dose-dependent effect on viral load in the brain, with titers being equivalent to control-treated mice at 20 and 2 µg of SKT05 (**Fig. 6G**). As expected, 100% of the mice administered 200 µg of SKT20 survived. In contrast, all mice provided 60, 20, or 2 µg of SKT20 lost weight and most met the criteria for euthanasia (**Fig. 6H-I**). As anticipated, SKT20 dose did not impact viral burden in the brain (**Fig. 6J**). Surprisingly, the dose needed for reduction of viral RNA in the periphery was much lower; a 2 µg dose of SKT05 and a 20 µg dose of SKT20 significantly reduced viral RNA in the spleen (**Fig. S4**). Importantly, a 10-fold reduction in SKT05 dose (20 µg) recapitulated the weight loss, survival, and brain viral burden of the 200 µg dose of SKT20 suggesting that functional avidity correlates with protection.

To separate the importance of epitope specificity from binding avidity for *in vivo* protection, we used SKT14, a broadly reactive anti-alphavirus mAb that competes with SKT05 for binding to VEE VLP but has an IC_50_ value (0.30 µg/ml) for VEEV pseudovirus neutralization, i.e., a range similar to SKT20 (0.27 µg/ml) (*11*). First, we determined the binding affinity to TC-83-infected cells and showed a binding curve similar to SKT20 with a 2.5-fold difference in EC_50_ value (0.68 µg/ml) compared to SKT05 (**Fig. 6A, D**). When SKT14 was administered one day before TC-83 infection, mice gained weight (**Fig. 6K**) and were protected from lethality (**Fig. 6L**). SKT14-treated mice had viral RNA levels in the brain that were between those of SKT05 and SKT20 (**Fig. 6M**). These results suggest that reduction in viral RNA in the brain may correlate better with antibody epitope. We next tested the Fc-dependency of SKT14 for VEEV protection. SKT14 LALA-PG treated mice all lost weight and succumbed to the TC-83 challenge (**Fig. 6N-O**). These results demonstrate that despite having a similar binding epitope to that of SKT05, SKT14 is dependent on Fc-engagement for protection suggesting that functional avidity is a better predictor for requirement of Fc effector functions.

To verify that avidity dictates Fc-dependency for *in vivo* efficacy, we administered a 10-fold higher dose of the SKT14 and SKT20 LALA-PG variants to compensate for differences in VEEV pseudovirus neutralization, as compared to SKT05. Mice received 200 µg of SKT14, 200 µg or 2 mg of SKT14 LALA-PG, or 2 mg of a control antibody in prophylaxis to lethal TC-83 challenge. As expected, mice administered 200 µg of SKT14 survived while mice provided 200 µg of SKT14 LALA-PG lost weight and succumbed to the infection (**Fig. 6P-Q**). However, mice administered 2 mg of SKT14 LALA-PG lost weight but had 100% survival rate (**Fig. 6P-Q**). Similar results were seen with SKT20 (**Fig. 6R-S**). Mice that received 2 mg of SKT20 LALA-PG survived with only minor weight loss (**Fig. 6R-S**). Despite the differing epitopes of SKT14 and SKT20, they both share Fc-dependent mechanisms that can be overcome by providing an equivalent potency dose as SKT05.

### SKT05 reduces clinical disease during therapeutic administration and protects against other alphaviruses with conditional requirement for Fc effector functions

Goals of mAb development against alphaviruses include therapeutic efficacy and breadth of protection against closely and distantly related viruses. Since SKT05 was superior in binding to infected cells, VEEV pseudovirus neutralization, and protection when administered in prophylaxis, as compared to SKT20, we evaluated the therapeutic potential and cross protection of SKT05. In these studies, we also assessed the necessity of Fc effector functions for protection. Following infection with TC-83, mice were administered 200 µg of SKT05 or SKT05 LALA-PG at 1, 2, or 3 dpi. We saw nearly complete survival of mice administered SKT05 at 1, 2, and 3 dpi with one of eight mice meeting end-point criteria in the 2 dpi administration group (**Fig. 7A**). Mice administered SKT05 LALA-PG at 1 and 2 dpi were protected from lethal challenge but only 20% survived in the group provided the mAb at 3 dpi (**Fig. 7A**). Mice receiving therapeutic doses later during infection had a marginal increase in weight loss (**Fig. 7B**). Treatment with the LALA-PG variant at 3 dpi resulted in significant weight loss and the survivors did not recover to starting weight by study end (**Fig. 7B, *right***).

**Figure 7.**
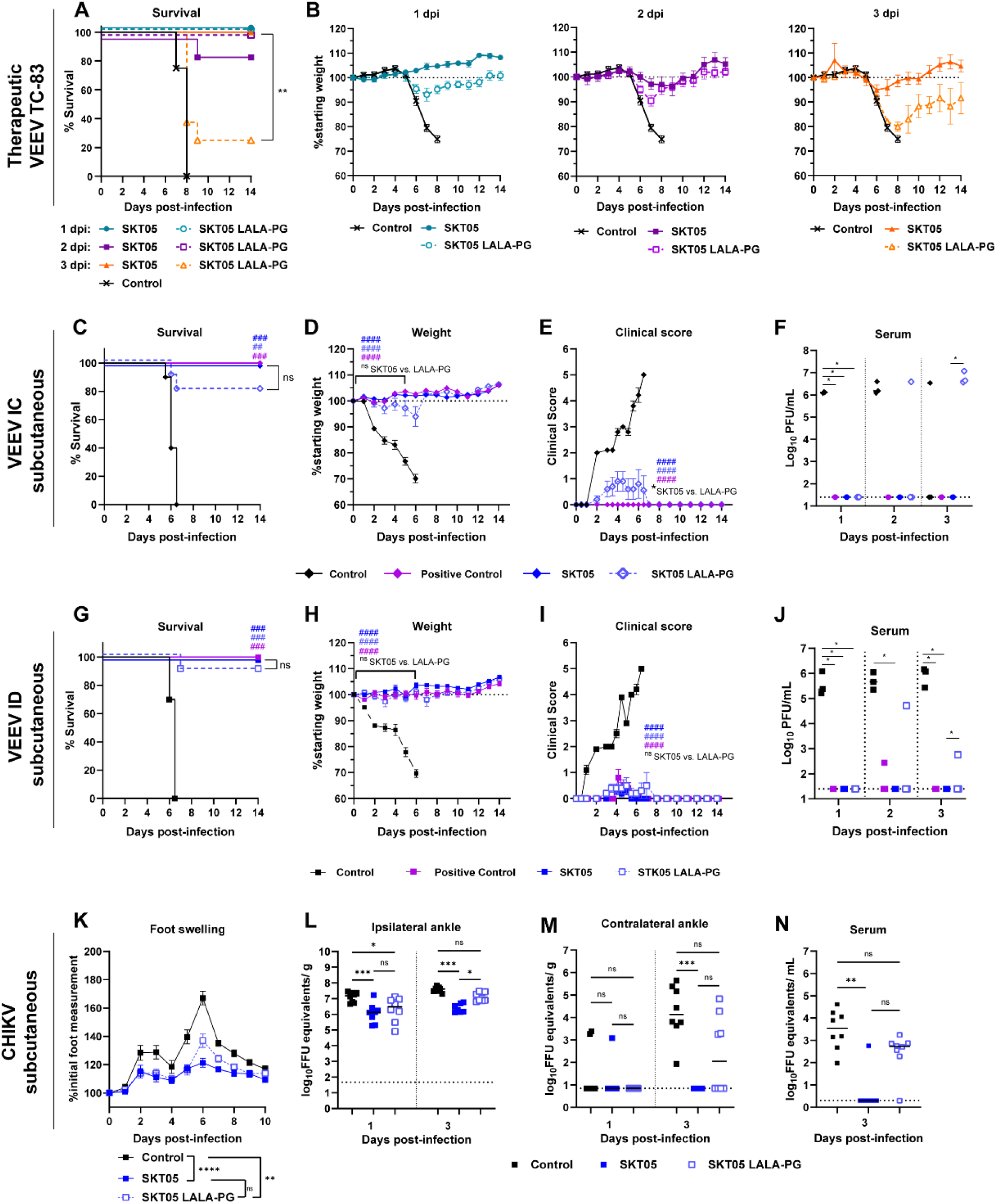
SKT05 reduces clinical disease independent of Fc effector functions during therapeutic administration and against other alphaviruses. (**A-B**) C3H/HeN mice were infected with TC-83 then administered 200 µg of SKT05, SKT05 LALA-PG, or isotype control antibody at 1, 2, or 3 dpi. Mice were followed for survival (A) and weight loss (B) for 14 days (n = 8/group; 2 independent experiments). Statistical significance was determined by a Log-rank test between SKT05 to SKT05 LALA-PG at each time of administration (A), one-way ANOVA with a Tukey’s post-test of AUC analysis from 0-7 dpi comparing all groups at each time point (B). No significant difference was observed for any comparison in (B). (**C-J**) Balb/C mice were treated with 200 µg of SKT05, SKT05 LALA-PG, a positive control mAb (1A3B7), or a control mAb at 1 h post-subcutaneous challenge with 10^3^ PFU of VEEV INH-9813 (IC subtype; C-F) or ZPC-738 (ID subtype; G-J). Mice were followed for 14 days for survival (C, G), weight loss (D, H), and clinical score (E, I) (n = 10/group; 2 independent experiments). Viremia was assessed on 1, 2, and 3 dpi (F, J) (n = 3/ group; 1 independent experiment). Statistical significance was determined by a Log-rank test between treatment groups to control (#; symbol color) and SKT05 to SKT05 LALA-PG (C and G), one-way ANOVA with a Sidak’s post-test of AUC analysis from 0-5 dpi (D-E) or 0-6 dpi (H-I) comparing treatment groups to control mAb (#; symbol color) and SKT05 to SKT05 LALA-PG, or Kruskal Wallis with a Dunn’s post-test between treatment groups to control and SKT05 to SKT05 LALA-PG at each time point (F and J). (**K-N**) C57BL/6J mice were treated with 200 µg of SKT05, SKT05 LALA-PG, or a control mAb 1 day prior to infection with 10^3^ FFU of CHIKV in the ipsilateral footpad. (K) Mice were monitored for foot swelling. Data shows the mean ± SEM of three independent experiments (n=14/group). Statistical significance was determined by one-way ANOVA with Tukey’s post-test of AUC analysis from all time points comparing all groups. (L-N) Additional mice were euthanized at 1 or 3 dpi and viral RNA load was determined by RT-qPCR (n = 8/group; 2 independent experiments). Data shows the median and statistical significance was determined by Kruskal-Wallis with a Dunn’s post-test comparing all groups. All weight loss and clinical score data represent mean ± SEM.

Next, we assessed protection of SKT05 against epizootic and enzootic strains of VEEV. Mice were inoculated by subcutaneous injection in the rear footpad with VEEV subtype IC or subtype ID, to mimic the natural route of transmission. At 1 h post infection, SKT05, SKT05 LALA-PG, a positive control mAb (1A3B-7), or control mAb was administered. For VEEV IC challenge, SKT05 treatment protected mice from lethality, weight loss, clinical disease, and viremia (**Fig. 7C-F).** Eight out of 10 mice survived with SKT05 LALA-PG administration: a result not statistically different from SKT05 administration (**Fig. 7C**). SKT05 LALA-PG-treated mice showed maximally 5% weight loss and minimal, but apparent, clinical disease compared to SKT05-treated mice (**Fig. 7D-E**). Viremic mice treated with SKT05 LALA-PG had viral loads similar to control mice at 2 and 3 dpi (**Fig. 7F**). For VEEV ID challenge, SKT05-treated mice were completely protected, and LALA-PG mice had 90% survival (**Fig. 7G**). SKT05 and LALA-PG groups lost minimal weight (**Fig. 7H**) and had average clinical scores less than 1 (**Fig. 7I**). Importantly, viremia was not detectable in SKT05-treated mice, while viremia was only observed in one mouse administered SKT05 LALA-PG at 2 and 3 dpi (**Figure 7J**).

We next wanted to determine if the Fc-independent mechanism of protection observed with VEEV extended to arthritogenic alphaviruses. SKT05, SKT05 LALA-PG, or a control mAb were administered one day prior to CHIKV infection. Mice were monitored for foot swelling and viral load was assessed at 1 and 3 dpi. SKT05 and SKT05 LALA-PG-treated mice had significantly reduced foot swelling compared to control mice (**Fig. 7K**). However, LALA-PG mice had notable increases in foot swelling at 6 dpi, which paralleled a peak in swelling observed in the control mice, although to a lesser extent (**Fig. 7K**). At 1 dpi, SKT05 and LALA-PG administration reduced viral load compared to control mice in the infected (ipsilateral) ankle **(Fig. 7L**). At 3 dpi, LALA-PG-treated mice had viral loads similar to control mice. A similar pattern was observed in disseminated tissues (**Fig. 7M and S5**). Analogous to the VEEV subcutaneous challenges, LALA-PG-treated mice, at 3 dpi, had detectable viremia (**Fig. 7N**). These results indicate that Fc effector functions are not required for early therapeutic clinical efficacy of SKT05 but are necessary for control of viral burden for both encephalitic and arthritogenic alphaviruses.

## DISCUSSION

Here, we defined the mechanism of protection for two broadly reactive anti-alphavirus mAbs, SKT05 and SKT20, during TC-83-induced encephalitis. SKT05 and SKT20 reduced inflammation in the brain and protected mice from VEEV infection in the absence of potent authentic VEEV neutralization. SKT05 strongly bound to the surface of infected cells and blocked viral egress in a neuronal cell line. This likely contributed to the Fc-independent survival observed with prophylaxis and/or therapeutic administration during infection with multiple VEEV subtypes or CHIKV. In contrast, SKT20 required Fc effector functions for protection. Functional avidity, as a correlate of binding to infected cells and potency of pseudovirus neutralization, rather than epitope, predicted dependence on Fc effector functions for survival. Indeed, the requirement for Fc mediated protection could be overcome by adjusting the mAb dose to compensate for the lower binding and pseudovirus neutralization potency.

Virus-specific and broadly reactive mAbs targeting E1 epitopes have been identified and characterized for multiple alphaviruses (*2*). Many broadly reactive E1-specific antibodies fail to or poorly neutralize authentic virus which is thought to be related to binding cryptic E1 epitopes that may only be exposed in certain stages of the viral life cycle (*2, 4, 30*). Consequentially, anti-E1 antibodies may alternatively engage with FcγRs for *in vivo* efficacy. In our study, SKT05 and SKT20 reduced early viral RNA in the brain. Using a single-cycle entry inhibition assay in Vero cells, we showed that SKT05 and SKT20 only mildly impacted TC-83 entry which is consistent with minimal neutralization using our standard FRNT assay. This lack of inhibition was not cell type dependent as the mAbs did not inhibit entry into the LUHMES.

Broadly reactive E1-targeting mAbs have previously been shown to inhibit viral egress in Vero and Neuro2a cells (*4, 7*). We observed egress inhibition in Vero cells with the most robust inhibition observed with SKT05 and SKV09 (5 – 10-fold reduction in virion release). Surprisingly, SKT05 inhibited TC-83 egress in LUHMES cells with greater magnitude (60-fold reduction) than in Vero cells. SKT20 inhibited egress with reduced potency relative to SKT05, paralleling the limited control of viral replication in the brain by this antibody. Since SKT05 did not require Fc effector functions to mediate survival or early control of virus replication, this data suggests that egress inhibition in certain types of neuronal cells may be the primary mechanism of protection for SKT05. Furthermore, the requirement for Fc effector functions can be overcome by dose escalation suggesting that egress inhibition may be a dominant mechanism of protection. However, the clinical outcome relies on the potency of mAb binding to infected cells.

SKT05 provided superior therapeutic efficacy following TC-83 challenge. Other broadly reactive anti-E1 mAbs, DC2.112, DC2.315 (*7*), EEEV-138, EEEV-346 (*4*), and 1A4B-6 (*30*), demonstrated >70% survival in mice when administered prior to VEEV-SINV, VEEV TrD, or EEEV (FL93-939) challenge. From the therapeutic perspective, DC2.112 and DC2.315, administered at 1 dpi, provided 100% protection in mice challenged with VEEV (ID) but only 20 and 50% protection when administered at 2 dpi, respectively (*7*). EEEV-138 and EEEV-346 resulted in 30% and 40% survival when administered 1 day following EEEV challenge (*4*). Notably, SKT05 supersedes the prophylactic and therapeutic potential of these antibodies, by providing 100% survival with a 20 µg (1 mg/kg) dose in prophylactic dose and ≥ 80% protection at 2 dpi and 3 dpi in TC-83-challenged mice. Furthermore, Fc effector functions were only required when SKT05 was administered at 3 dpi, while Fc independent protection for DC2.112 was lost when administered 1 dpi. Other VEEV-specific mAbs have been shown to protect in mice and non-human primates at 2 dpi (*31–33*). However, the necessity of Fc engagement has not been well established. More recently, an E2-binding, humanized VEEV-specific mAb, h5F, was shown to require Fc effector functions when administered at 2 dpi following TC-83 challenge (*9*). While not addressed in our study, we hypothesize that a combination therapy may extend the therapeutic window beyond 3 dpi. Additionally, the therapeutic efficacy of SKT05 against arthritogenic alphavirus challenge remains unclear. Previous work with anti-CHIKV mAbs suggests that Fc effector functions would be required for therapeutic protection (*7, 8*). However, the therapeutic window for Fc-dependent protection needs further investigation. Overall, these results emphasize the potential for prophylactic and therapeutic use of SKT05 during alphavirus infections and warrant future studies evaluating SKT05 efficacy in non-human primate models toward a clinical application.

Unlike SKT05, SKT20 required Fc effector functions for protection at our standard (200 µg) dose. SKT20 reduced viral RNA in the brain at 6 dpi, presumably through FcγR-mediated clearance, which may have contributed to survival. Based on previous studies with CHIKV and Mayaro virus (MAYV) (*7, 8, 10*), virus was cleared through Fc-FcγR interactions on monocytes. FcγR interaction on monocyte or macrophage could increase cellular activation, as observed with SKT20 treatment. However, SKT20 administration reduced the number of Ly6C^hi^ monocytes compared to SKT20 LALA-PG treatment, which is supported by reduction in the monocyte chemoattractant, CCL2, and molecules produced by activated monocytes such as CCL4, CCL5, IL-6, and IL-1β. An earlier study demonstrated increased Ly6C^+^ monocytes in TC-83 infected mice as early as 2 dpi in the olfactory bulb and significant levels in the cortex until 6 dpi (*14*). It is unresolved whether monocyte recruitment into the brain is protective or pathogenic during VEEV encephalitis and it could be that the timing of infiltration is critical. Since we only assess a single time-point, it is possible that SKT20 modulates early chemokine response and influx of monocytes to reduce inflammation. Additional studies are needed to address timing and spatial distribution of monocyte recruitment.

T cells are recruited to the brain starting around 6 dpi and aid in clearance of VEEV (*14, 34, 35*). The proportion of CD4^+^ T cells in SKT20 mice is increased compared to SKT20 LALA-PG treatment, but the percentage of CD4^+^ and CD8^+^ T cells expressing CD69 is reduced. This may be related to differences in T cell activation. SKT20 controls viral replication in the periphery, which could reduce the amount of antigen available for T cell activation. In the subcutaneous VEEV models, SKT05 LALA-PG administration failed to control viral replication in the periphery past 3 dpi. While peripheral viral loads were not assessed with SKT20 LALA-PG administration, it is likely the same viral kinetic shift would be present and thus have provided similar antigen levels as the control treated mice for T cell stimulation. Alternatively, fundamental studies conducted with neuroadapted SINV showed that host-generated antibodies act synergistically with interferon-γ produced by T cells to control SINV infection through noncytolytic clearance of neurons (*36–38*). However, this mechanism is primarily observed with antibodies targeting E2 and does not require an Fc domain (*39, 40*).

In this study, we established a direct relationship between functional avidity, as determined by mAb pseudovirus neutralization and infected cell binding, and the requirement of Fc effector functions *in vivo*. *In vitro* antibody neutralization is typically used as a correlate for *in vivo* protection. However, our studies suggest that pseudovirus neutralization and cell surface binding are just as important of predictors. The pseudovirus particles only contain the alphavirus envelope glycoproteins. Without capsid, the pseudovirus particles would likely be less structured compared to an authentic alphavirus and may more closely resemble the E2/E1 trimers present on the surface of infected cells, potentially providing analogous readouts. The ability to overcome the requirement of Fc effector functions for SKT20 and SKT14 protection by equalizing the dose relative to SKT05 functional avidity indicates that mAbs are not restricted to one mechanism for protection. Higher avidity would drive clustering of mAbs on the cell surface and promote antibody dependent cellular cytotoxicity, antibody dependent cellular phagocytosis, and complement dependent cytotoxicity (*41*). However, the same idea of increased mAb clustering could be connected to enhanced egress inhibition. If early viral control in this model dictates clinical outcome, enhanced egress inhibition may be sufficient to prevent lethality. Importantly, assessing mAb avidity could accelerate the identification of broadly protective mAbs that are more effective against emerging and re-emerging alphaviruses.

## LIMITATIONS

The epizootic strains of VEEV (subtypes IAB and IC) are select agents and require high containment facilities. For logistical reasons, we used the VEEV BSL2 strain TC-83 (IAB) to evaluate the mechanisms of protection for SKT05 and SKT20, then performed confirmatory experiments with virulent strains of VEEV and showed complementary results. Furthermore, we did not address the therapeutic window for SKT05 against virulent strains of VEEV. This would need to be evaluated in future studies. Finally, we assumed that SKT14 and SKT05 target the same epitope based on competition ELISA data; however, structural analysis needs to be performed to confirm similar epitope recognition and binding angle.

## Materials and Methods

### Study Design

The goals of this study were to determine the mechanisms of protection for two broadly reactive anti-alphavirus antibodies using mouse models of alphavirus infection and relate functional mAb features to the requirement for Fc effector functions for protection. *In vivo* studies were conducted in mouse models that are well-established for alphaviruses and the number of mice used for each study, to ensure the studies were appropriately powered, was determined based on historical experiments and known variation in the results. Male and female mice were used for the VEEV TC-83 and CHIKV mouse studies and female mice were used for VEEV TrD, INH-9813, and ZPC-738 studies. Predetermined endpoints were used for tissue collection. For survival studies, mice were humanely euthanized once a pain score of 3 or weight loss criteria was met. Number of independent experiments and experimental replicates are detailed in each figure legend. All data points were used for analysis; no outliers were excluded. To remove bias, histological samples were blinded prior to processing and scoring by a board-certified veterinary pathologist.

### Cells

African Green Monkey Kidney Cells (Vero; CCL-81) and Baby Hamster Kidney cells (BHK-21; CCL-10) were obtained from American Type Culture Collection (ATCC). Vero and BHK cells were cultured at 37°C with 5% CO_2_ in complete media [Dulbecco’s Modified Essential Medium (DMEM) supplemented with 5% heat-inactivated fetal bovine serum (HI-FBS; Omega) and 10 mM HEPES (Gibco)].

Lund Human Mesencephalic Cells (LUHMES) were obtained from Applied Biological Materials Inc. (Cat. T0284). Cell culture flasks and dishes were sequentially coated with 50µg/ml poly-l-ornithine hydrobromide (Sigma) overnight at RT followed by 1µg/ml fibronectin (Sigma) for 6 hr at 37°C, rinsed with ddH2O, and allowed to fully air dry at room temperature before cell plating. LUHMES were maintained in proliferation media [DMEM:F12 (Sigma-Aldrich) containing 1% N2 supplement (ThermoFisher Sci.), 1X Penicillin-Streptomycin solution (Corning), 2mM L-glutamine (Corning)] with 40ng/mL recombinant human Fibroblast Growth Factor (FGF-basic, Peprotech) added right before use. Cells were plated at 100,000 cells/well in 24 well dishes. Media was changed 24 h after plating and the cells were allowed to grow for 2 more days before using for experiments.

### Viruses

The pVE/IC-92 cDNA clone encoding the full-length VEEV (strain TC-83) genome was acquired from the World Reference Center for Emerging Viruses and Arboviruses at The University of Texas Medical Branch, rescued as previously described (Kinney et al., 1998), and passaged once on Vero cells. TC-83 stocks were sequenced to confirm the absence of the A3G mutation. VEEV INH-9813 (IC strain) stock was passaged three times on Vero cells, VEEV Trinidad donkey (TrD; IAB strain) stock was received from DynPort Vaccine Company (DVC) and prepared by Commonwealth Biotechnologies Inc., and VEEV ZPC-738 (ID strain) stock was passaged once on BHK cells. Virus titer was determined by standard plaque assay on Vero76 cells. CHIKV (strain AF15561) was generated from a cDNA clone as previously described (*42*) and passaged once on BHK cells. Viral stocks were titrated by focus forming assay (FFA; TC-83 and CHIKV) or plaque assay (VEEV TrD, INH-9813, and ZPC-738), as previously described (*6, 43*). All experiments with VEEV strains INH-9813, TrD, and ZPC-783 and CHIKV were conducted in BSL-3/ABSL-3 conditions.

### Antibodies

Macaque mAbs, SKT05, SKT20, SKV09, and ITS103.01 (anti-SIV control mAb) were generated as rhesus macaque IgG1, as previously described (Sutton et.al, 2023). The LALA-PG (L234A L235A P329G) mutation was inserted into the heavy chain plasmids. The paired heavy and light chain plasmids were co-transfected to Expi293 cells by Expifectamine 293 transfection kit following the manufacturer’s instructions. Full length IgG was purified by rProtein A Sepharose Fast Flow antibody purification resin. SKT05, SKT20, and their LALA-PG variants all include the half-life extending LS mutation.

### Mouse studies

Experiments related to TC-83 and CHIKV challenges were carried out in accordance with the recommendations in the Guide for the Care and Use of Laboratory Animals of the National Institutes of Health in compliance with the National Institute of Allergy and Infectious Diseases (NIAID) Animal Care and Use Committee (ACUC) under the approved protocol LVD 6E. Experiments related to VEEV TrD, INH-9813, and ZPC-738 were conducted under an Institutional Animal Care and Use Committee (IACUC) approved protocol at USAMRIID in compliance with the Animal Welfare Act, PHS Policy, and other Federal statutes and regulations relating to animals and experiments involving animals. The facility where this research was conducted is accredited by the Association for Assessment and Accreditation of Laboratory Animal Care International and adheres to principles stated in The Guide for the Care and Use of Laboratory Animals, National Research Council, 2011.

#### VEEV TC-83 challenge

C3H/HeN purchased from Charles River Laboratories were used between 6-8 weeks of age in equal ratio of male to female mice per experiment. Antibodies were diluted in PBS solution and administered by intraperitoneal injection (i.p.). For therapeutic administrations of antibody, mice were briefly sedated using isoflurane prior to administration of antibodies. Mice were challenged intranasally either pre- or post-antibody administration with 10^7^ FFU of TC-83 virus in PBS (40 uL total; 20 uL per nare) under anesthesia with 2,2,2-Tribromoethanol (Avertin; Fisher Scientific). For weight-loss and challenge studies, mice were weighed daily and humanely euthanized when ≥25% of their starting weight was lost or they reached a pain score of 3. For tissues collections, mice were euthanized and perfused with PBS prior to tissue harvest.

#### VEEV IAB (TrD), IC (INH-9813), or ID (ZPC-738) challenge

Six-to eight-week-old, specific pathogen-free, female BALB/c mice from Charles River were used at ABSL-3. Mice were exposed to target dose of 10^3^ PFU of VEEV IAB strain via aerosol. The aerosol challenge was generated using a Collison Nebulizer to produce a highly respirable aerosol (flow rate 7.5 ± 0.1 L/minute). The system generates a target aerosol of 1 to 3 µm mass median aerodynamic diameter determined by aerodynamic particle sizer. For VEEV IC and ID challenges, mice were infected subcutaneously with 10^3^ PFU in the rear footpad. Mice received a single dose of 200 μg of antibody intraperitoneally 1 hour post virus exposure. On days 1, 2, and 3 post-exposure, blood was collected from 3 animals/group for use in plaque assay to determine viremia. Clinical observations were performed daily for signs of disease and weight loss. Once mice reached a clinical score ≥3, observations were increased to twice daily. Mice that displayed severe signs of disease were humanely euthanized. Clinical scores were based on the below scale:

1 = reduced grooming – minor alteration in fur or soiled coat

2 = ruffled fur – severe alteration in fur; raised fur with ruffled appearance

3 = hunched posture – outward curvature of the spine at the back resulting in a hunch

4 = lethargic – decreased activity; animal not moving around as much as normal

5 = neurological signs (circling/hind limb paralysis) or unresponsive when stimulated – animal does not respond or move even when provoked

#### CHIKV challenge

All CHIKV challenge studies were conducted at ABSL-3. C57BL/6J mice were purchased from Jackson Laboratory and used at 4-weeks of age in equal numbers of males and females. Mice were administered SKT05, SKT05 LALA-PG, or a control antibody (ITS103.01) (in PBS) by i.p. injection one day prior to subcutaneous inoculation in the rear footpad with 10^3^ FFU of CHIKV in Hanks Balanced Salt Solution (HBSS) supplemented with 1% HI-FBS under isoflurane anesthesia. Swelling of the ipsilateral foot was measured (width x height) prior to infection and for 10 days following infection using digital calipers. Other mice were euthanized at 1 and 3 dpi, perfused with PBS, and the indicated tissues were harvested for viral burden analysis.

### RNA extraction and RT-qPCR

Perfused tissues were homogenized in 1 mL of viral infection medium (DMEM supplemented with 2% HI-FBS, 10 mM HEPES, and 100 U/mL of penicillin and streptomycin) using Zirconia/Silica beads in a MagNA Lyser for 60s at 6,000 rpm. For viral load analysis, homogenates were clarified by centrifugation at 10,000 rpm for 5 min. For TC-83 and CHIKV-infected tissues, RNA was extracted from the clarified homogenate using the Kingfisher Duo Prime with MagMax-96 Viral RNA isolation kit (ThermoFisher) or RNeasy kit (Qiagen), respectively, following the manufacturer’s instructions. To determine viral burden, equal quantities of RNA were added to Taqman fast virus 1-step master mix (ThermoFisher) with TC-83 nsP3 specific primers/probes (Forward: 5’-CCATATACTGCAGGGACAAGAA-3’, Reverse: 5’-CACTGAAGAGTCGTCGGATATG-3’,Probe:5’-56’FAM/ATGACTCTC/ZEN/AAGGAAGCAGTGGCT/3IABkFQ/-3’) or CHIKV E1 specific primers/probes (Forward: 5’-TCGACGCGCCATCTTTAA-3’, Reverse: 5’-ATCGAATGCACCGCACACT-3’,Probe:5’-/56 FAM/ACCAGCCTG/ZEN/CACCCACTCCTCAGAC/3IABkFQ/-3’) (*44*). Reactions were run on a QuantStudio 3-Real-Time PCR System and viral RNA isolated from TC-83 and CHIKV viral stocks were used to generate a standard curve based on FFU equivalents. All tissues were normalized to gram of tissue or mL of serum.

For host gene-expression analysis, 30 mg weight/volume of brain tissue homogenate was mixed with 1:10 volume of TRIzol™ (ThermoFisher) and subjected to phenol: chloroform phase separation per the manufacturer’s instructions. Isolated RNA was quantitated by Nanodrop and equally added to Taqman RNA-to-CT™ 1-Step Kit (ThermoFisher,) master mix for RT-qPCR with the Taqman assay primer/probes for mouse gene-transcripts for *Icam-1* (Mm00516023_m1), *Mmp-9* (Mm00442991_m1), *Cxcl9* (Mm00434946_m1), *Cxcl10* (Mm00445235_m1), and *Gapdh* (Mm99999915_g1).

### Cytokine and chemokine analysis

Perfused brains were collected at 5 or 6 dpi and homogenized in viral infection medium as described above. Following homogenization, samples were mixed with a 1x protease inhibitor solution containing a cOmplete™, Mini EDTA-Free Protease Inhibitor Cocktail to prevent degradation of respective analytes. Samples were stored at -80°C until further use. Cytokines and chemokines from 5 dpi samples (SKT05, SKT20, and Control) were analyzed for respective analytes using a Bio-Plex Pro Mouse Cytokine 31-Plex Assay kit (Bio-Rad). Samples collected at 6 dpi (SKT20, SKT20 LALA-PG, and Control) were analyzed using a Bio-Plex Pro Mouse Chemokine 23-Plex assay kit (Bio-Rad) following the manufacturer’s instructions. Each experiment included naïve brain homogenates for determination of baseline cytokine or chemokine concentrations for respective Bio-plex assay kits.

### Flow Cytometry

Following euthanasia and perfusion, brains were collected at 6 dpi and stored on ice in HBSS prior to downstream processing. Brain tissue was minced then incubated in an enzymatic digestion solution [RPMI (Gibco) supplemented with 2.5 mg/mL of Type IV Collagenase (Thermofisher), 100 µg/mL of Liberase TL (Sigma), 10 µg/mL DNase I (Sigma), and 15 mM HEPES] at 37°C on a plate rocker for 20 minutes. Tissue was sporadically agitated by pipetting throughout the 20-minute incubation. Cells were filtered through a 70-µm cell strainer, pelleted, and resuspended in 70% Percoll. Cells were isolated from myelin and other debris with a 30-37-70% Percoll gradient. Pellets were washed with cold 1x HBSS and resuspended in fluorescence-activated cell sorting (FACS) buffer (1% FBS in PBS). Single cell suspensions were counted and compensation controls were made with a pool of single cell suspensions from both infected and mock infected animals. Cells were blocked for FcγR binding (BioLegend clone 93; 1:50), and surface stained using fluorochrome-conjugated anti-mouse antibodies: CD45 BUV395 (clone 30-F11; BD Biosciences; 1:200), CD11B FITC (clone M1/70; BioLegend; 1:200), CD19 BUV737 (clone 1D3; BD Biosciences; 1:200) CD3 PerCP-Cy5.5. (clone KT3.1.1; BioLegend; 1:100), CD4 BV605 (clone RM4-5; BioLegend; 1:100), CD8 APC (clone 53.6.7; BioLegend; 1:100), NK1.1 PE-Cy7 (clone PK136;BioLegend; 1:200), Ly6G APC-Cy7 (clone 1A8; BioLegend; 1:200), Ly6C BV650 (clone HK1.4; BioLegend; 1:400), F4/80 BV421 (clone T45-2342; BD Biosciences; 1:100), MHCII BV711 (clone MF/114.15.2; BioLegend; 1:400), and CD69 PE (clone S15049F; BioLegend; 1:100). Cell viability was determined by exclusion of fixable viability dye (Aqua) (Thermofisher). Samples were run on a BD LSRFortessa flow cytometer and analyzed using FlowJo version 10.10 (Flojo, LLC).

### Egress inhibition assay

One day prior to infection, 1.0 x 10^5^ Vero cells were seeded into 24-well plates in complete medium. On the day of the experiment, extra wells were sacrificed to determine cell count. Cells were washed 1x with PBS then infected at an MOI 1 in viral infection medium for 1 h at 37°C. Following infection, cells were washed 4x with PBS then mAbs diluted to 10 µg/mL, 1 µg/mL, and 0.1 µg/mL in egress medium (DMEM supplemented with 2% FBS, 10 mM HEPES, and 25 mM NH_4_Cl to prevent *de novo* infection) was applied to cells for 6 h at 37°C with 5% CO_2_. Supernatant was harvested at 1 and 6 hpi and subjected to RNA extraction as previously described above. Egress assays were completed similarly with LUHMES except proliferation media was used for infections and proliferation media supplemented with NH_4_Cl for subsequent incubation with mAbs.

### Histology and in situ hybridization of viral RNA

Mice were provided SKT05, SKT05 LALA-PG or a control antibody 1 day prior to intranasal challenge with TC-83. At 1 and 6 dpi, mice were euthanized and perfused with PBS. Mice were then perfused with 4% paraformaldehyde (PFA) in PBS for total fixation. After fixation, the heads were scalped, the calvarium were removed, and the skulls were placed in 4% PFA at a minimum ratio of 1:10 (tissue volume: 4% PFA volume), at room temperature for 24 h. Each skull was rinsed with PBS and water followed by decalcification with 14% EDTA on a plate rocker, at room temperature, for approximately 14 days. The EDTA solution was replaced initially after the first 24 h and then replaced every 3 days during the period of decalcification. The skulls were rinsed with water 3x and stored in wetted gauze with 10% neutral buffered formalin until further processing. Tissues were divided using a midsagittal cut, dehydrated with increasing concentrations of ethanol, and paraffin embedded. Sequential sections of one side of the tissue were stained with H&E or probed for VEEV RNA using RNAscope2.5 VS Universal AP reagent kit [Advanced Cell Diagnostics, Inc. (ACD)] with a VEEV nsp3 specific probe (ACD, Cat. 404509). Representative images were acquired with an Olympus BX51 microscope and Olympus DP80 camera using cellSens software (Olympus). All tissues were evaluated and scored blind by a board-certified veterinary pathologist as described in the supplementary material and methods.

### mAb binding to the surface of live infected cells

Vero cells were seeded at 1.0 X 10^6^ cells/well in 6-well plates one day prior to infection. On the day of infection, wells were sacrificed for counting. Vero cells were infected at an MOI of 1 for 1 h in viral infection medium at 37°C. Following infection, viral inoculum was removed, cells were washed 2x, and media was replaced with viral infection medium. At 18 hpi, cells were trypsinized, washed, counted, and equally split into wells of a 96-well U-bottom plate to ensure a minimum of 80,000 cells/well would be stained. Cells were stained with serial dilutions of mAb in FACS buffer for 1 h at 4°C. After washing, goat anti-human AF647-conjugated IgG was applied in FACS buffer and cells were stained for 1 h at 4°C. Following washing, cells were fixed in 4% PFA in PBS for 10 min at 4°C. Cells were washed and resuspended in FACS buffer and samples were run on a BD LSRFortessa flow cytometer and analyzed using FlowJo version 10.10 (Flojo, LLC).

### Statistical analysis

Statistical analysis was performed using GraphPad Prism Version 10. The statistical test and multiple comparisons post-test, when applicable, used for each analysis is described here or in the figure legend. The appropriate analysis was determined based on number of groups being compared, variation, points at or above the limit of detection for the assay, and normalization of the data. For area under the curve (AUC) analysis, only time points where all mice were alive were included in the analysis. The day range for each AUC analysis is included in the figure legends. The Kaplan-Meier curves with more than one comparison were corrected for multiple comparisons. Unless otherwise noted in the figure legend: *, p< 0.05; **, p< 0.01, ***, p < 0.001, ****, p < 0.0001, and ns, not significant.

## List of Supplementary Materials

Material and Methods

Fig. S1 to S5

Table S1 to S2

## Acknowledgements

We like to thank the NIAID comparative medicine branch for animal care and technical assistance with the TC-83 and CHIKV studies. We like to thank the technical support of Ashley Piper, Yvonne Gonzalez-Nederstigt, and Lauren Panny in completion of the VEEV TrD, IC and ID studies and the USAMRIID Aerobiology, Animal Clinical Pathology, Telemetry (AAT) team for their assistance with the TrD aerosol exposure.

## Author contributions

Conceptualization: VC, MSS, MR, JMF

Methodology: VC, MSS, CLG, JLV, JPS, DL, TMK, CWB, MR, JMF

Investigation: VC, MSS, CLG, DK, MMD, MG, CG, TYC, CSC, JMF

Visualization: VC, MSS, CLG, CSC

Funding acquisition: TMK, CLG, CWB, MR, JMF

Supervision: TMK, CWB, MR, JMF

Writing – original draft: VC, CSC, JMF

Writing – review & editing: VC, MSS, DK, MMD, MG, TYC, DL, JLV, TMK, CSC, CWB, MR, JMF

## Disclosures

The opinions, interpretations, conclusions, and recommendations presented are those of the author and are not necessarily endorsed by the U.S. Army or Department of Defense.

#Contractor – this does not constitute an endorsement by the U.S. Government of this or any other contractor.

## Data and materials availability

All data supporting these results will be available in source data files within this paper or on a public repository. Additional requests should be directed to the corresponding author. A material transfer agreement may be required for any shared reagent.

## REFERENCES

1. J. Jose, J. E. Snyder, R. J. Kuhn, A structural and functional perspective of alphavirus replication and assembly. Future Microbiol 4, 837–856 (2009).

2. A. S. Kim, M. S. Diamond, A molecular understanding of alphavirus entry and antibody protection. Nat Rev Microbiol 21, 396–407 (2023).

3. S. Raju et al., A chikungunya virus-like particle vaccine induces broadly neutralizing and protective antibodies against alphaviruses in humans. Sci Transl Med 15, eade8273 (2023).

4. L. E. Williamson et al., Therapeutic alphavirus cross-reactive E1 human antibodies inhibit viral egress. Cell 184, 4430–4446 e4422 (2021).

5. L. A. Powell et al., Human mAbs Broadly Protect against Arthritogenic Alphaviruses by Recognizing Conserved Elements of the Mxra8 Receptor-Binding Site. Cell Host Microbe 28, 699–711 e697 (2020).

6. J. M. Fox et al., Broadly Neutralizing Alphavirus Antibodies Bind an Epitope on E2 and Inhibit Entry and Egress. Cell 163, 1095–1107 (2015).

7. A. S. Kim et al., Pan-protective anti-alphavirus human antibodies target a conserved E1 protein epitope. Cell 184, 4414–4429 e4419 (2021).

8. J. M. Fox, et al., Optimal therapeutic activity of monoclonal antibodies against chikungunya virus requires Fc-FcgammaR interaction on monocytes. Sci Immunol 4, (2019).

9. J. L. Schwedler et al., Therapeutic efficacy of a potent anti-Venezuelan equine encephalitis virus antibody is contingent on Fc effector function. MAbs 16, 2297451 (2024).

10. J. T. Earnest et al., Neutralizing antibodies against Mayaro virus require Fc effector functions for protective activity. J Exp Med 216, 2282–2301 (2019).

11. M. S. Sutton et al., Vaccine elicitation and structural basis for antibody protection against alphaviruses. Cell 186, 2672–2689 e2625 (2023).

12. K. E. Steele et al., Comparative neurovirulence and tissue tropism of wild-type and attenuated strains of Venezuelan equine encephalitis virus administered by aerosol in C3H/HeN and BALB/c mice. Vet Pathol 35, 386–397 (1998).

13. J. G. Julander et al., C3H/HeN mouse model for the evaluation of antiviral agents for the treatment of Venezuelan equine encephalitis virus infection. Antiviral Res 78, 230–241 (2008).

14. M. D. Cain et al., Virus entry and replication in the brain precedes blood-brain barrier disruption during intranasal alphavirus infection. J Neuroimmunol 308, 118–130 (2017).

15. E. P. Williams et al., Deep spatial profiling of Venezuelan equine encephalitis virus reveals increased genetic diversity amidst neuroinflammation and cell death during brain infection. J Virol 97, e0082723 (2023).

16. A. Schafer, C. B. Brooke, A. C. Whitmore, R. E. Johnston, The role of the blood-brain barrier during Venezuelan equine encephalitis virus infection. J Virol 85, 10682–10690 (2011).

17. A. Sharma, M. Bhomia, S. P. Honnold, R. K. Maheshwari, Role of adhesion molecules and inflammation in Venezuelan equine encephalitis virus infected mouse brain. Virol J 8, 197 (2011).

18. S. Paessler et al., Alpha-beta T cells provide protection against lethal encephalitis in the murine model of VEEV infection. Virology 367, 307–323 (2007).

19. J. B. Dietrich, The adhesion molecule ICAM-1 and its regulation in relation with the blood-brain barrier. J Neuroimmunol 128, 58–68 (2002).

20. B. S. Hollidge et al., Toll-like receptor 4 mediates blood-brain barrier permeability and disease in C3H mice during Venezuelan equine encephalitis virus infection. Virulence 12, 430–443 (2021).

21. A. Rani, S. Ergun, S. Karnati, H. C. Jha, Understanding the link between neurotropic viruses, BBB permeability, and MS pathogenesis. J Neurovirol 30, 22–38 (2024).

22. B. A. Schoneboom, K. M. Catlin, A. M. Marty, F. B. Grieder, Inflammation is a component of neurodegeneration in response to Venezuelan equine encephalitis virus infection in mice. J Neuroimmunol 109, 132–146 (2000).

23. A. L. Phelps, et al., Tumour Necrosis Factor-alpha, Chemokines, and Leukocyte Infiltrate Are Biomarkers for Pathology in the Brains of Venezuelan Equine Encephalitis (VEEV)-Infected Mice. Viruses 15, (2023).

24. J. T. Earnest et al., The mechanistic basis of protection by non-neutralizing anti-alphavirus antibodies. Cell Rep 35, 108962 (2021).

25. T. Rulker et al., Isolation and characterisation of a human-like antibody fragment (scFv) that inactivates VEEV in vitro and in vivo. PLoS One 7, e37242 (2012).

26. M. Hezareh, A. J. Hessell, R. C. Jensen, J. G. van de Winkel, P. W. Parren, Effector function activities of a panel of mutants of a broadly neutralizing antibody against human immunodeficiency virus type 1. J Virol 75, 12161–12168 (2001).

27. M. D. Cain et al., Post-exposure intranasal IFNalpha suppresses replication and neuroinvasion of Venezuelan Equine Encephalitis virus within olfactory sensory neurons. J Neuroinflammation 21, 24 (2024).

28. N. M. Kafai et al., Entry receptor LDLRAD3 is required for Venezuelan equine encephalitis virus peripheral infection and neurotropism leading to pathogenesis in mice. Cell Rep 42, 112946 (2023).

29. K. E. Steele, N. A. Twenhafel, REVIEW PAPER: pathology of animal models of alphavirus encephalitis. Vet Pathol 47, 790–805 (2010).

30. A. E. Calvert et al., Exposing cryptic epitopes on the Venezuelan equine encephalitis virus E1 glycoprotein prior to treatment with alphavirus cross-reactive monoclonal antibody allows blockage of replication early in infection. Virology 565, 13–21 (2022).

31. C. W. Burke et al., Therapeutic monoclonal antibody treatment protects nonhuman primates from severe Venezuelan equine encephalitis virus disease after aerosol exposure. PLoS Pathog 15, e1008157 (2019).

32. L. M. O’Brien, S. A. Goodchild, R. J. Phillpotts, S. D. Perkins, A humanised murine monoclonal antibody protects mice from Venezuelan equine encephalitis virus, Everglades virus and Mucambo virus when administered up to 48 h after airborne challenge. Virology 426, 100–105 (2012).

33. S. A. Goodchild et al., A humanised murine monoclonal antibody with broad serogroup specificity protects mice from challenge with Venezuelan equine encephalitis virus. Antiviral Res 90, 1–8 (2011).

34. C. B. Brooke, D. J. Deming, A. C. Whitmore, L. J. White, R. E. Johnston, T cells facilitate recovery from Venezuelan equine encephalitis virus-induced encephalomyelitis in the absence of antibody. J Virol 84, 4556–4568 (2010).

35. N. E. Yun et al., CD4+ T cells provide protection against acute lethal encephalitis caused by Venezuelan equine encephalitis virus. Vaccine 27, 4064–4073 (2009).

36. R. Burdeinick-Kerr, J. Wind, D. E. Griffin, Synergistic roles of antibody and interferon in noncytolytic clearance of Sindbis virus from different regions of the central nervous system. J Virol 81, 5628–5636 (2007).

37. D. E. Griffin, Recovery from viral encephalomyelitis: immune-mediated noncytolytic virus clearance from neurons. Immunol Res 47, 123–133 (2010).

38. G. K. Binder, D. E. Griffin, Interferon-gamma-mediated site-specific clearance of alphavirus from CNS neurons. Science 293, 303–306 (2001).

39. B. Levine et al., Antibody-mediated clearance of alphavirus infection from neurons. Science 254, 856–860 (1991).

40. S. Ubol, B. Levine, S. H. Lee, N. S. Greenspan, D. E. Griffin, Roles of immunoglobulin valency and the heavy-chain constant domain in antibody-mediated downregulation of Sindbis virus replication in persistently infected neurons. J Virol 69, 1990–1993 (1995).

41. S. C. Oostindie, G. A. Lazar, J. Schuurman, P. Parren, Avidity in antibody effector functions and biotherapeutic drug design. Nat Rev Drug Discov 21, 715–735 (2022).

42. A. W. Ashbrook et al., Residue 82 of the Chikungunya virus E2 attachment protein modulates viral dissemination and arthritis in mice. J Virol 88, 12180–12192 (2014).

43. P. Pal et al., Development of a highly protective combination monoclonal antibody therapy against Chikungunya virus. PLoS Pathog 9, e1003312 (2013).

44. A. Bakovic et al., Venezuelan Equine Encephalitis Virus nsP3 Phosphorylation Can Be Mediated by IKKbeta Kinase Activity and Abrogation of Phosphorylation Inhibits Negative-Strand Synthesis. Viruses 12, (2020).

