## Supplemental Material for "Requirement for Fc effector function is overcome by binding potency for broadly reactive anti-alphavirus antibodies"

**Supplemental Files**

**Supplemental materials and methods**

**Focus forming assay (FFA)**

Tissues were weighed, homogenized, and clarified in infection medium as described in the “RNA extraction and RT-qPCR” section of the material and methods. Vero cells, were infected with serial dilutions of clarified homogenates for 1.5 hours at 37°C. The inoculum was removed, and cells were overlayed with a 1% methylcellulose (Sigma-Aldrich) in Minimum Essential Medium (MEM, Sigma-Aldrich) supplemented with penicillin and streptomycin, 10 mM HEPES, and 2% HI-FBS. Cells were fixed 16 h later with 4% paraformaldehyde (PFA; Electron Microscopy Sciences) in PBS. Cells were washed with PBS, permeabilized with perm buffer (PBS supplemented with 0.1% saponin and 0.1% BSA) and stained overnight at 4°C with 500 ng/mL of VEEV-specific macaque antibody SKV09 diluted in perm buffer. The next day cells were washed with wash buffer (PBS supplemented with 0.05% Tween-20), then incubated with an anti-monkey IgG conjugated to HRP diluted in perm buffer (1:5000; KPL) for 2 h. Cells were washed and foci were visualized using True-Blue Peroxidase substrate (SeraCare) and counted using an Immunospot plate reader (CTL).

**Plaque assay for VEEV IAB, IC, and ID viremia**

Serial dilutions of sera were made in MEM with 2% HI-FBS, 1% HEPES, and 2% Pen/Strep. Six-well plates of Vero76 cells were infected with 0.2 mL of each serial dilution per well in duplicate wells and plates were incubated at 37°C for 1 h. After 1 h incubation, cells were overlaid with 0.6% agarose in Basal Medium Eagle (BME) with 10% HI-FBS, and 2% Pen/Strep, and incubated overnight at 37°C, 5% CO_2_. A second overlay containing 0.6% agarose in BME with 10% HI-FBS, 2% Pen/Strep, and 4% of total volume neutral red vital stain was added to the wells, and plates were further incubated for 18-28 h for visualization of plaques. Plaques were counted following incubation with stain overlay. Virus stock was utilized as a positive control and media only was used as a negative control. Limit of detection of the assay is 25 PFU/mL.

**Neutralization of VEEV Env-pseudotyped lentiviral vectors/reporters**

Neutralization assays were similarly performed as described previously (*11*). Briefly, one day before infection 10^4^ cells/well of 293A cells in Minimum Essential Media (MEM) supplemented with 5% FBS and 1% penicillin-streptomycin (M5 media) were seeded into tissue culture treated black & white 96-well isoplate. Titrated amounts of pseudotyped lentiviral reporters (with p24 levels of about 50 ng/mL) were incubated with 4-fold serially diluted mAbs from 50 mg/mL or 10 mg/mL at 37°C in a 5% CO_2_ incubator for 45 min. Next, 50mL/well of the virus:mAb solution were added to 293A cells and incubated at 37°C in a 5% CO_2_ incubator for 2 h. After incubation, 100 mL/well of M5 media was added to the plate. An anti-SIV mAb served as a negative control. After 24 h of incubation, cells were lysed using cell lysis buffer. Luciferase activity was measured using the Micro-Beta JET after incubation with a luciferase assay reagent according to the manufacturer’s protocol. All experiments were performed in triplicate. Inhibition values were calculated as follows: inhibition (%) = [1 – [luciferase activity (cps) in pseudotyped lentiviral reporter–infected cells incubated with the indicated dilutions of mAb]/[luciferase activity (cps) in pseudotyped lentiviral reporter–infected cells incubated with the same dilutions of control mAb] X 100. Calculations for the 50, 80, and 90% inhibitory concentrations were made with GraphPad Prism.

**Focus reduction neutralization test (FRNT)**

mAbs were serially diluted in viral infection medium and then incubated with 10^2^ FFU of TC-83 for 1 h at 37°C. Virus-antibody complexes were then added to Vero cells and incubated for 1.5 h at 37°C. Cells were overlaid with Minimum essential medium (MEM) and 1% methylcellulose supplemented with 2% HI-FBS, 10 mM HEPES, and penicillin and streptomycin and incubated for 16 h. The cells were fixed with 1% PFA/PBS for 1 h and then washed and permeabilized in perm buffer (PBS supplemented with 0.1% BSA and 0.1% saponin). The cells were stained with 500 ng/mL of VEEV-specific macaque antibody SKV09 diluted in perm buffer, washed with wash buffer (PBS supplemented with 0.05% Tween-20), then incubated with an anti-monkey IgG conjugated to HRP diluted in perm buffer (1:5000; KPL). Cells were washed then foci were visualized using True-Blue Peroxidase substrate (SeraCare) and counted using an Immunospot plate reader (CTL). The number of foci with mAbs were compared to wells with no mAb to determine percent relative infection. Non-linear regression with the top and bottom constrained to 100 and 0, respectively, was used to determine the IC_50_ value (GraphPad Prism).

**Quantification of macaque IgG in tissues**

Serum and perfused, whole brain samples were collected at 5 dpi from mice treated with respective mAbs and infected with TC-83. Next, serum was heat inactivated for 30 min at 56°C while whole brains were homogenized, clarified, and then heat inactivated at 56°C for 1 hr. Following inactivation, 500 uL of clarified homogenate was concentrated 10-fold with Corning® Spin-X® UF Centrifugal Concentrator (VWR). Unlabeled goat anti-human IgG Fc-specific and cross-absorbed against mouse IgG (Sigma) was absorbed overnight at 4°C on Maxisorp immunocapture ELISA plates (Thermo Fisher Scientific) in sodium bicarbonate buffer (pH 9.3). The next day, plates were washed and then blocked with 2% bovine serum albumin (BSA) in PBS (blocking buffer) at 37°C for 1 h. Plates were washed, serum and clarified brain homogenate were serially diluted in blocking buffer and added to the wells for 1 h at 4°C. Plates were washed and then incubated for 1 h at 4°C with biotin-conjugated goat anti-human H&L (Jackson ImmunoResearch Laboratories). Following washing, plates were incubated with streptavidin-HRP for 30 min at room temperature. Plates were developed with TMB-One Step Substrate, the reaction was stopped with H_2_SO_4,_ and absorbance was acquired at 450 nm. Standard curves were run in parallel and analyzed using nonlinear regression to determine the concentration of the macaque monoclonal antibodies in the brain and serum of mice.

**Human FcγRI ELISA**

Recombinant human Fc gamma receptor 1 (hFcgRI/CD64) (R&D Systems) diluted to 2 µg/mL in sodium bicarbonate buffer (pH 9.3) was absorbed overnight at 4°C on Maxisorp immunocapture ELISA plates (Thermo Fisher Scientific). The next day, plates were washed with ELISA wash buffer then blocked in blocking buffer for 1 h at 37°C. Following blocking, SKT05, SKT20, SKT05 LALA-PG, and SKT20 LALA-PG were applied to plates in blocking buffer at 10 µg/mL, 1 µg/mL, and 0.1 µg/mL for 2 h at room temperature. Plates were washed and peroxidase labeled, goat anti-human IgG (H+L) (Seracare) was applied at 1:2000 dilution in blocking buffer for 1 h at room temperature. Following washing, the plate was developed with TMB One-Step Substrate (Thermo Fisher), and the reaction was neutralized with 2 M H_2_SO_4_ and absorbance readings were acquired at 450 nm using a BioTek Synergy H1 microplate reader.

**Single cycle entry inhibition assay**

Vero cells were seeded at 1.5 X 10^5^ cells/well in 24-well plates one day to the assay in complete media. LUHMES were seeded at 1.0 X 10^5^ cells/well in coated 24-well plates up to 3 days prior to the assay in proliferation media. On the day of the assay, wells were sacrificed for counting to ensure multiplicity of infection (MOI) of 0.5 was achieved for Vero and an MOI of 0.1 was achieved for LUHMES cell infections. These MOIs were predetermined to achieve 5-10% relative infection in cells by flow cytometry. TC-83 was diluted in viral infection medium (or proliferation media for LUHMES) and mixed 1:1 with mAbs diluted in 3-fold serial dilution starting at 10 µg/mL and incubated at 37°C for 1 h. Virus-antibody complexes and cell culture plates were then chilled at 4°C for 15 minutes. Virus-antibody complexes were added to chilled cells and plates were incubated on ice to permit attachment. After 1 h, virus-mAb complexes were removed, cells were washed 4x with PBS, then incubated for an additional 4 h at 37°C in the presence of viral infection medium (or proliferation media for LUHMES) supplemented with 25 mM NH_4_Cl. Cells were trypsinized, washed with PBS with 10 mM EDTA solution, and fixed for 20 minutes using BD Cytofix/Cytoperm™ Fixation/Permeabilization kit (BD Sciences). Following fixation, cells were permeabilized with the BD Perm/Wash Buffer and stained with 10 µg/mL of SKV09 in perm/wash buffer. After washing, goat anti-human AF647-conjugated IgG (Thermo Fisher) was added in perm/wash buffer for 1 h at 4°C. Following washing with perm/wash buffer, cells were resuspended in FACS buffer and run on a BD LSRFortessa flow cytometer and analyzed using FlowJo version 10.10 (Flojo, LLC).

**Supplemental Figures**

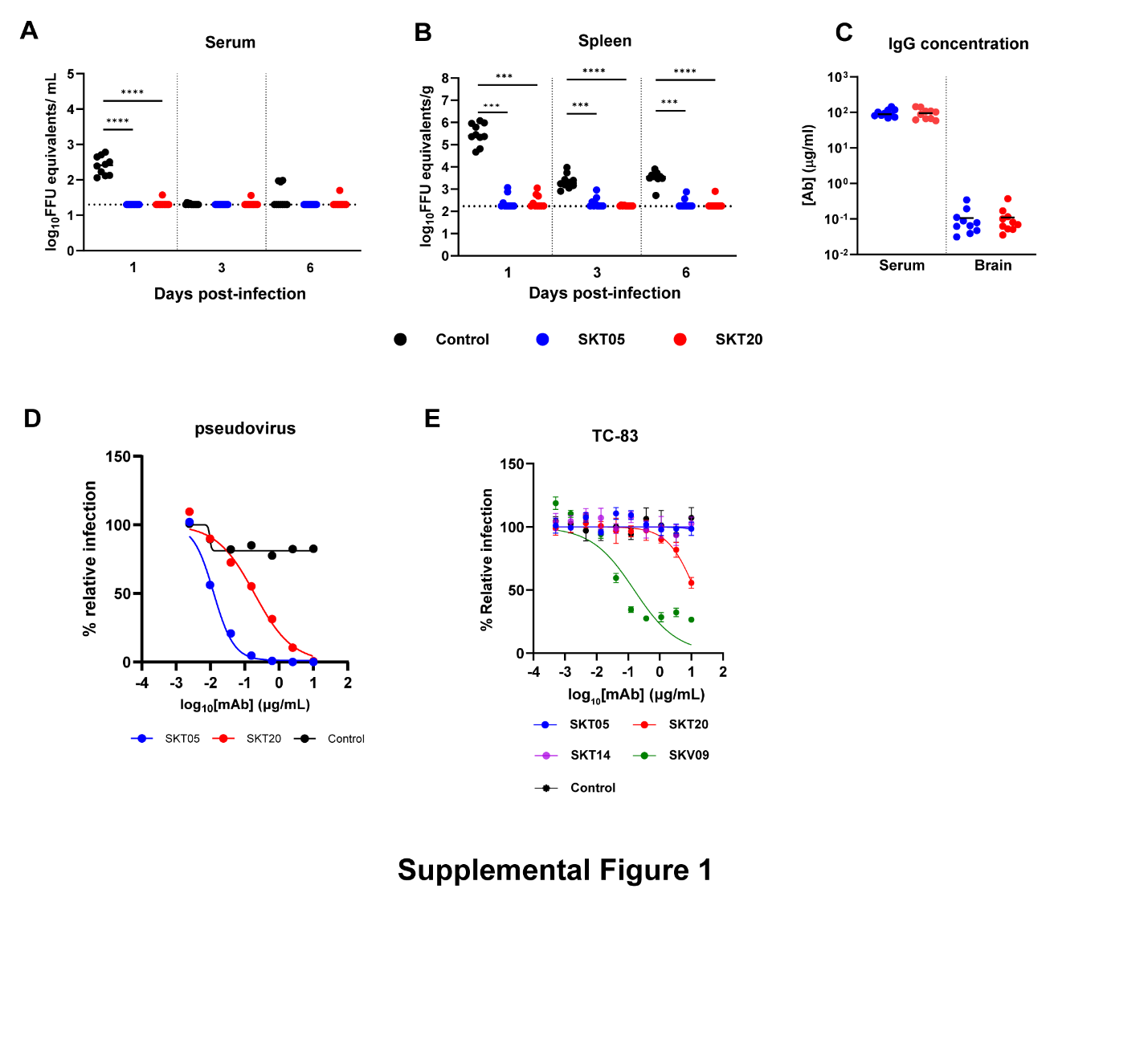

**Supplemental Figure 1. SKT05 and SKT20 demonstrate differential neutralization capacity *in vitro* and reduce viral burden in peripheral tissues.** (**A-B**) C3H/HeN mice were administered 200 µg of an isotype control, SKT05, or SKT20 1 day prior to inoculation with TC-83. At 1, 3, and 6 dpi, serum (A) and spleens (B) were collected to determine viral RNA load by RT-qPCR (n = 10/group; 2 independent experiments). The bars represent the median and statistical significance was determined by Kruskal-Wallis with a Dunn’s post-test comparing each group the isotype control. (**C**) The quantity of SKT05 or SKT20 in the serum or brain of mice was determined at 5 dpi by ELISA (n = 10/ group; two independent experiments). The bar represents the median. (**D**) Neutralization against VEEV Env-pseudotyped lentiviral reporter viruses. Data is representative of one independent experiment conducted in triplicate. (**E**) Focus reduction neutralization test (FRNT) with indicated mAbs and TC-83 is representative of two independent experiments conducted in duplicate. The graph represents the mean ± SD from two-independent experiments performed in duplicate. Non-linear regression with the top and bottom constrained to 100 and 0, respectively, was used to determine the IC_50_ value for neutralization curves.

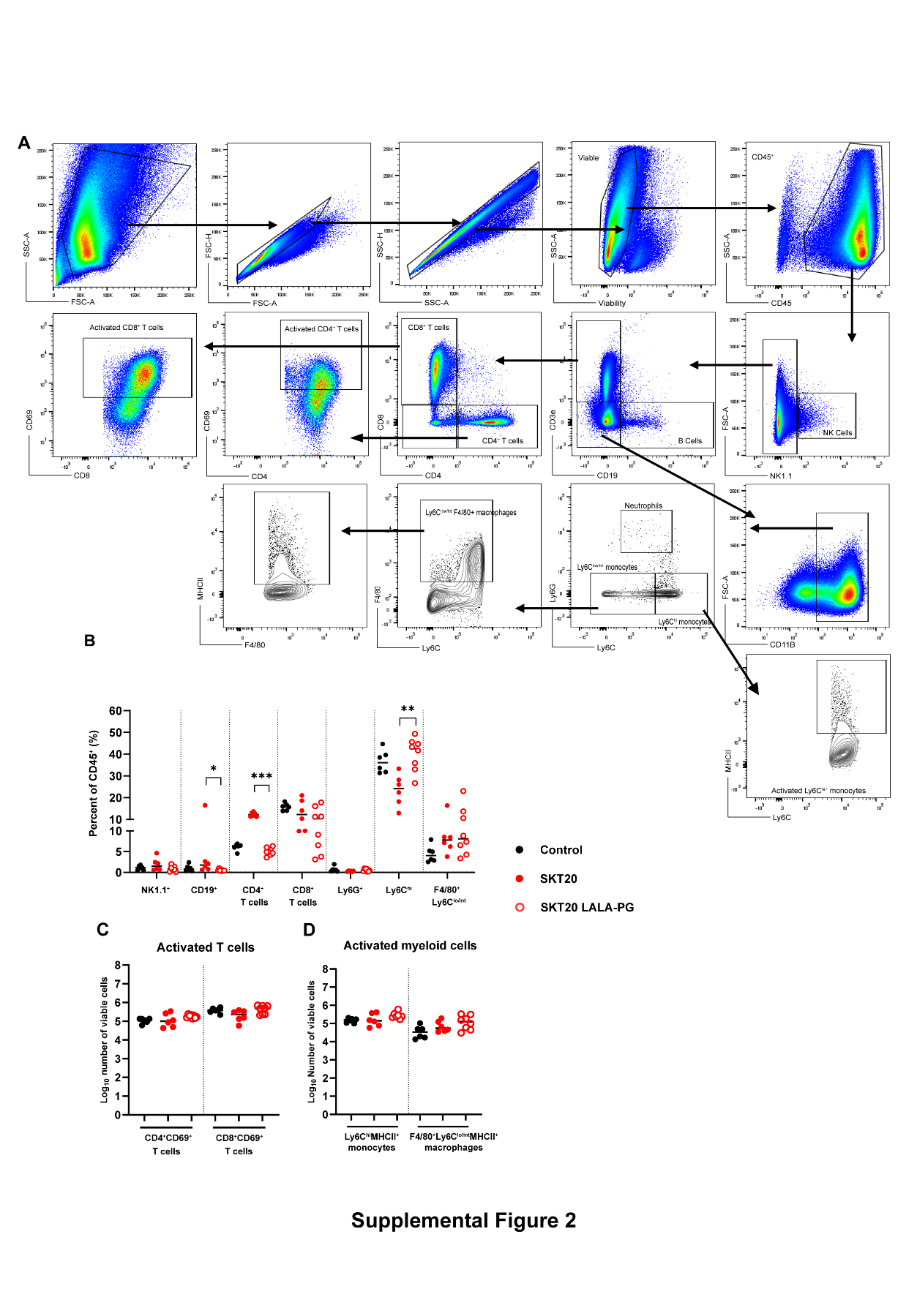

**Supplemental Figure 2. Gating scheme for infiltrating immune cells.**

(**A**) Gating scheme for identification of immune cell subsets. (**B-D**) C3H/HeN mice were given isotype control, SKT20, or SKT20 LALA-PG (200 µg) 1 day prior to infection with TC-83. At 6 dpi, brains were collected and digested. Single cell suspensions were stained and flow cytometry was performed to assess the percentage of NK cells (NK1.1^+^), B cells (CD19^+^), CD4^+^ T Cells, CD8^+^ T cells, neutrophils (Ly6G^+^), macrophages (Ly6C^lo/mid^ F4/80^+^), and monocytes (Ly6C^hi^), that comprised the CD45+ cell population (**B**). The number of activated (CD69^+^) CD4^+^ and CD8^+^ T cells (**C**) and activated (MHCII^+^) monocytes (Ly6C^hi^) and macrophages (Ly6C^lo/mid^ F4/80^+^) (**D**) was determined by flow cytometry. Bars indicate the median (n = 6 - 8/group; 2 independent experiments; Kruskal-Wallis with a Dunn’s post-test comparing all groups.

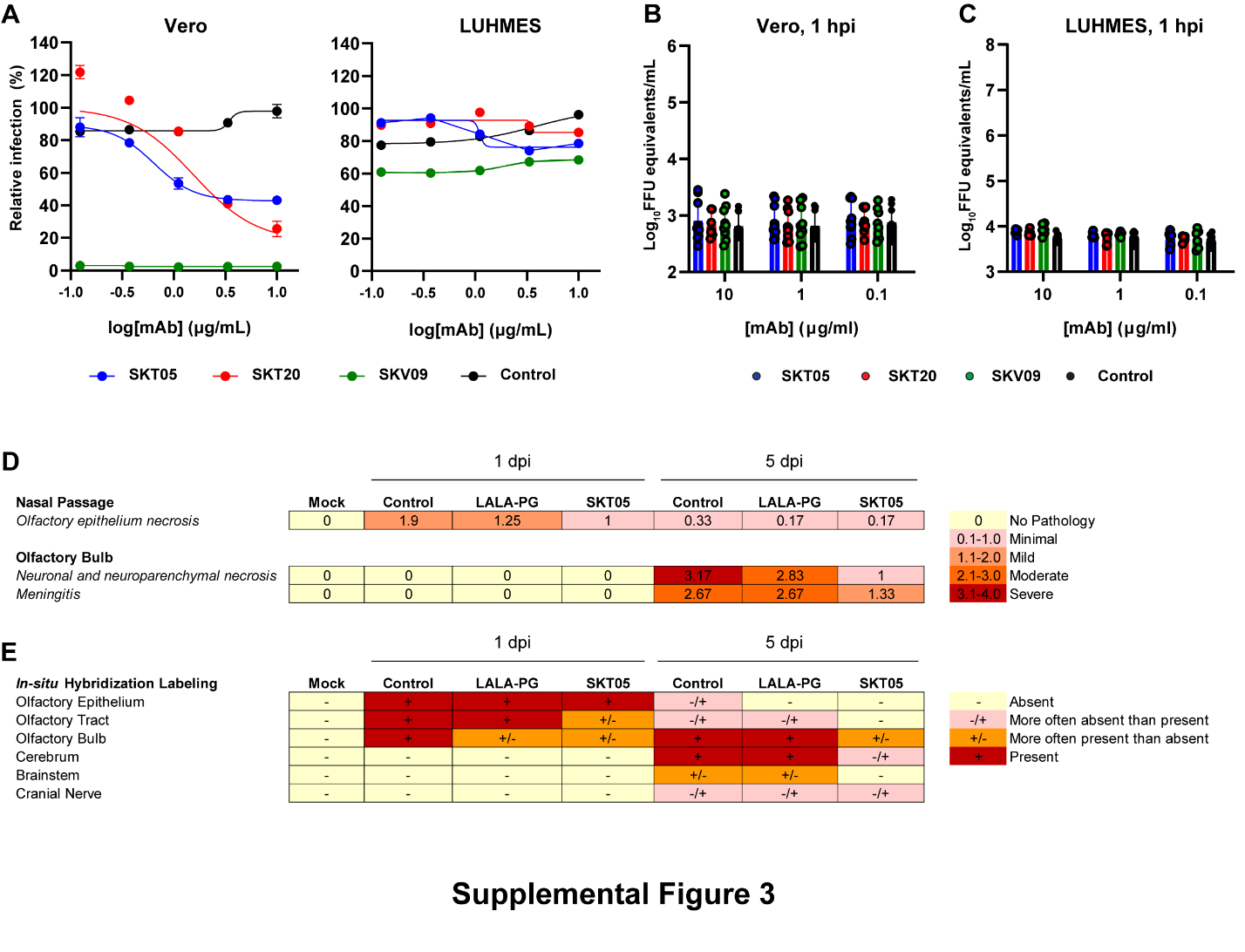

**Supplemental Figure 3. Single-cycle entry inhibition, viral egress, and immunohistochemical scoring of infected brains.** (**A**) A single-cycle, viral entry inhibition assay was performed on Vero cells or LUHMES. At 4 hpi cells were collected, fixed, stained for viral surface antigen, and flow cytometry was performed to determine the percent of infected cells. The IC_50_ was determined by non-linear regression. The data shows the represented mean ± SD of two independent experiments conducted in duplicate. Viral egress inhibition of SKT05, SKT20, SKV09, or an isotype control was evaluated in Vero cells **(B)** and LUHMES **(C).** Supernatants were collected at 1 hpi to quantify viral RNA by RT-qPCR. Data is representative of the mean ± SD of 2-3 independent experiments performed in triplicate. Statistical significance was determined by two-way ANOVA with Dunnett’s post-test comparing all groups to the isotype control. (**D-E**) C3H/HeN mice were pre-treated with 200 µg of SKT05, SKT05-LALA-PG, or an isotype control antibody one day before infection with TC-83. At 1 and 5 dpi, skulls with brains intact were harvested, fixed, followed by decalcification before paraffin embedding and sectioning. Slides containing midsagittal skull and brain sections were stained for VEEV RNA by *in situ* hybridization (ISH) or with hematoxylin and eosin (H&E) from animals harvested at 1 or 5 dpi. Slides were blinded prior to scoring of disease features such as olfactory epithelial necrosis, neuronal and parenchymal necrosis, and meningitis (**D**) or for presence or absence of viral RNA (**E**). (**D**) The average score is shown from two independent experiments (n = 6 - 8/group).

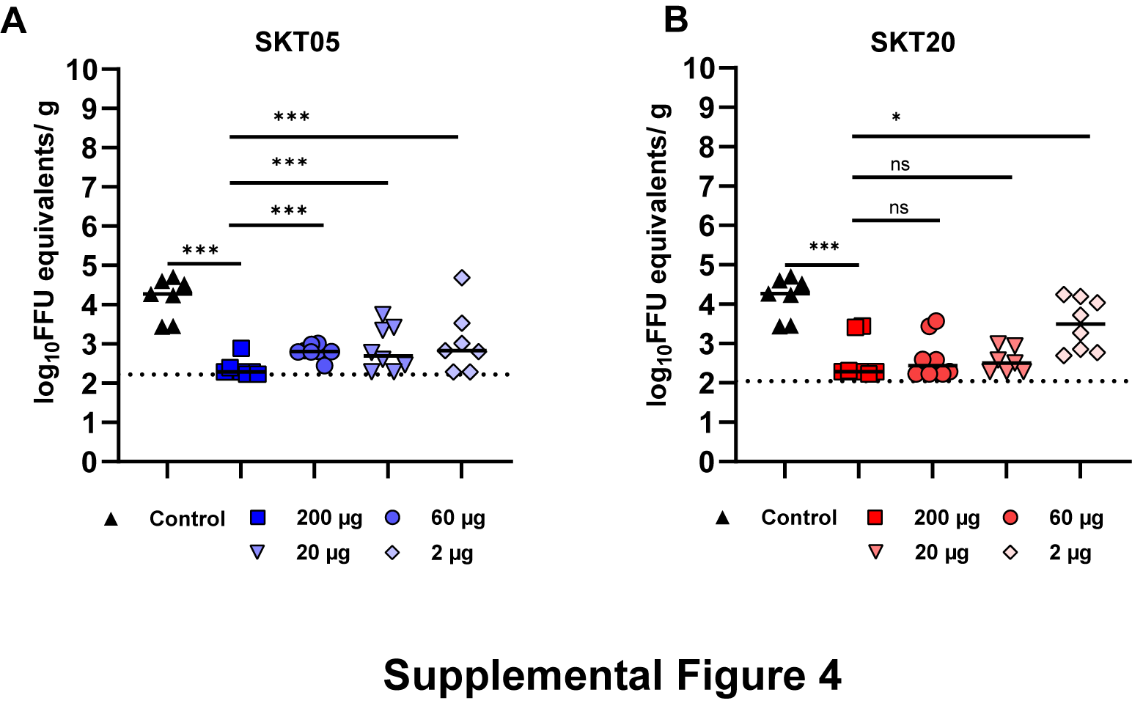

**Supplemental Figure 4. SKT05 and SKT20 dose-dependent protection is observed in peripheral tissues.** C3H/HeN mice were given 200, 60, 20, or 2 µg of SKT05, SKT20, or 200 µg of an isotype control 1 day prior to infection with TC-83. Viral loads were determined in the spleens at 5 dpi (n= 8/group; 2 independent experiments). Statistical significance was determined by Kruskal-Wallis with a Dunn’s post-test comparing each group the 200 µg dose. Bars represent the median.

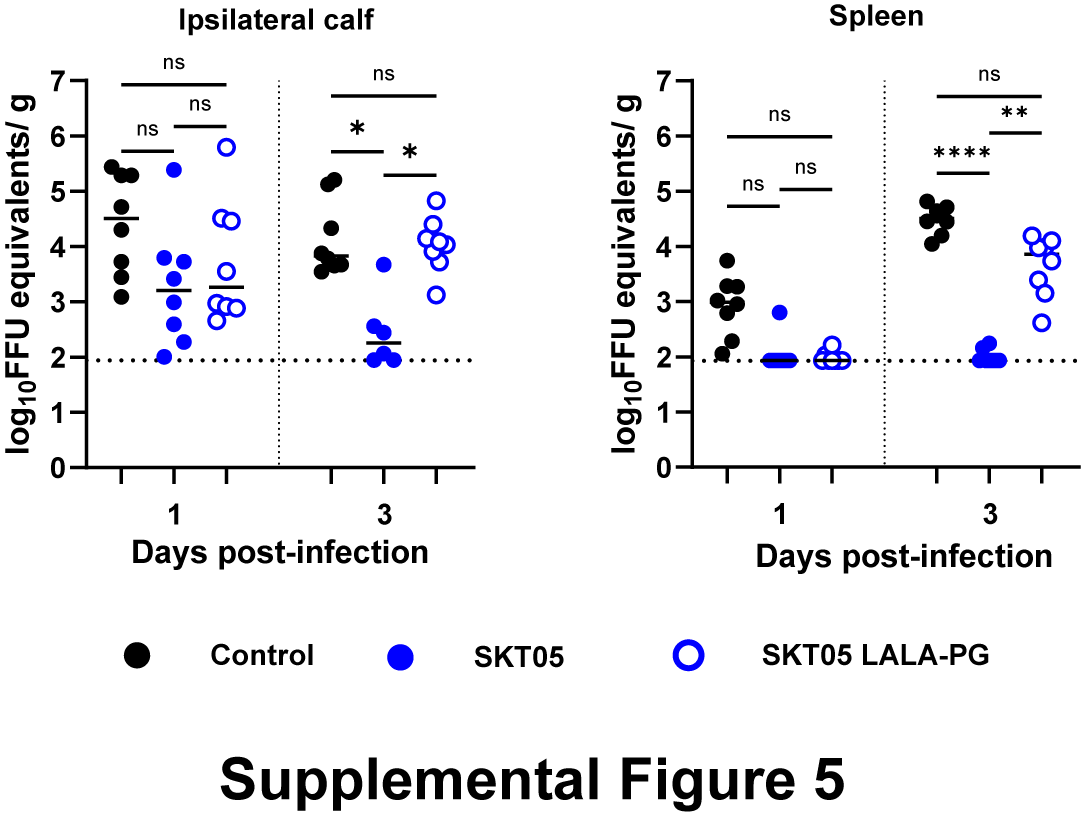

**Supplemental Figure 5. SKT05 Fc effector functions are required to maintain control of CHIKV in disseminated tissues.** C57BL/6J mice were given 200 µg of SKT05, SKT05 LALA-PG, or an isotype control 1 day prior to infection with CHIKV in the ipsilateral footpad. Mice were euthanized at 1 or 3 dpi and viral loads were determined in the ipsilateral calf and spleen (n = 8/group; 2 independent experiments). Bars represent the median and statistical significance was determined by Kruskal-Wallis with a Dunn’s post-test comparing all groups.

| **Median (pg/mL) (95% CI)^b^** | | | | ***P*-value^c^** | | |
| --- | --- | --- | --- | --- | --- | --- |
| **Analyte** | Control | SKT05 | SKT20 | SKT05 v. Control | SKT20 v. Control | SKT05 v. SKT20 |
| BCA-1 | 22,108 (21,537-23,308) | 6,014 (1,077-8,767) | 23,094 (22,741-25,027) | **0.008** | 0.35 | **<0.0001** |
| CCL12 | 4,955 (4,659-5,600) | 185.8 (47.7 - 295.6) | 2,292 (1,884-2,761) | **<0.0001** | **0.03** | **0.03** |
| CCL19 | 6,939 (6,139-8,724) | 918.2 (277.2-1,515) | 5,846 (4,837-6,750) | **<0.0001** | 0.19 | **0.01** |
| CCL2 | 52,441 (48,380-55,846) | 114.8 (74.83-146.5) | 5,682 (4,788-12,653) | **<0.0001** | **0.03** | **0.03** |
| CCL20 | 51.3 (42.6-54.6) | 36.0 (26.6-40.9) | 46.6 (40.9-51.4) | **0.0005** | >0.9999 | **0.0127** |
| CCL22 | 201 (165.3 - 210.3) | 41.1 (25.1 - 53.2) | 233.8 (167.4 - 268.1) | **<0.0001** | 0.003 | **>0.9999** |
| CCL3 | 3,989 (2810-5012) | 21.2 (15.4-25.3) | 571.1 (447.1-922.6) | **0.03** | **<0.0001** | **0.03** |
| CCL4 | 3,484 (2,902-5,362) | 19.81 (19.8-LOD) | 656.4 (422.0-952.9) | **<0.0001** | **0.03** | **0.03** |
| CCL5 | 10,370 (8,083-11,360) | 101.8 (24.20-161.8) | 3,147 (2,761-4,009) | **<0.0001** | **0.03** | **0.03** |
| CCL7 | 3,354 (3,241-3,398) | 32.09 (10.60-82.05) | 2,974 (2,899-3,198) | **<0.0001** | **0.03** | **0.03** |
| CTACK | 2,645 (2,369-2,852) | 1,796 (1,284-2,007) | 2,656 (2,235-3,148) | **0.0005** | >0.9999 | **0.0004** |
| CX3CL1 | 566.1 (470.0-733.2) | 420.1 (384.9-475.4) | 431.8 (405.0-482.0) | **0.002** | **0.008** | >0.9999 |
| CXCL1 | 980.0 (859.1-1,600) | LOD^d^ | 372.2 (341.8-749.7) | **<0.0001** | **0.02** | **0.04** |
| CXCL10 | 126,255 (114,957-137,495) | 10,990 (9,402-12,333) | 93,801 (83,679-102,571) | **<0.0001** | 0.06 | **0.02** |
| CXCl11 | 49,189 (44,240-55,387) | LOD | 24,419 (22,355-35,007) | **<0.0001** | **0.03** | **0.03** |
| CXCL12 | 1,316 (1,292-1,433) | 1,240 (1,086 -1,318) | 1,308 (1,243-1,423) | **0.01** | >0.9999 | 0.06 |
| CXCL16 | 1,101 (825.8-1,251) | 210.8 (152.7-240.3) | 555.5 (475.5-632.2) | **<0.0001** | 0.05 | **0.03** |
| CXCL17 | 257.1 (223.1-285.0) | 204.3 (176.4-237.2) | 264.4 (240.7-316.1) | **0.01** | >0.9999 | **0.0008** |
| ENA-78 | 535.4 (512.1-536.9) | 467.9 (376.7-543.1) | 467.9 (416.3-561.5) | **0.02** | 0.17 | >0.9999 |
| eotaxin | 323.9 (277.2-339.4) | 54.57 (46.1-61.9) | 170.6 (151.7-276.2) | **<0.0001** | 0.08 | **0.02** |
| eotaxin-2 | 14,086 (12,669-18,035) | 20,670 (15,853-21,903) | 1,1577 (9,386-15,199) | **0.04** | 0.76 | **0.001** |
| GM-CSF | 2.6 (1.6-3.1) | 0.1 (0.07-0.6) | 2.2 (1.5-2.6) | **<0.0001** | >0.9999 | **0.002** |
| I-309 | 56.4 (39.6-75.4) | 23.2 (18.9-25.3) | 51.9 (41.1-63.2) | **0.0002** | >0.9999 | **0.001** |
| IFNγ | 520.5 (319.3-840.5) | 8.2 (6.3-10.4) | 232.4 (201.7-532.2) | **<0.0001** | 0.20 | **0.01** |
| IL-10 | 427.5 (61.7-123.3) | 104.0 (61.7-147.8) | 340.7 (307.2-391.6) | **<0.0001** | 0.13 | **0.01** |
| IL-16 | 455.2 (407.5 - 567.5) | 208.9 (171.9-251.5) | 388.6 (376.2-437.6) | **<0.0001** | 0.49 | **0.006** |
| IL-1B | 32.3 (30.0-43.4) | 7.8 (4.6-10.4) | 27.6 (19.3-36.9) | **<0.0001** | 0.58 | **0.005** |
| IL-2 | 3.9 (3.4-4.1) | 2.9 (2.3-3.3) | 3.6 (2.9-4.1) | **0.002** | 0.98 | **0.04** |
| IL-4 | 31.0 (26.9-34.0) | 4.0 (10.4-21.1) | 26.9 (25.8-33.0) | **<0.0001** | 0.73 | **0.004** |
| Il-6 | 930.8 (702.1-1,069) | 3.4 (2.9-5.4) | 98.6 (64.7-344.4) | **<0.0001** | **0.03** | **0.04** |
| TNFα | 104.7 (46.3-187.6) | LOD | LOD | **0.0003** | **0.0003** | >0.9999 |

**Table S1: Proinflammatory cytokine and chemokine concentrations in brain tissue homogenates^a^**

^a^Mice were administered SKT05, SKT20, or an isotype control one day prior to intranasal infection with TC-83. At 5 dpi, brains were collected, homogenized, and analyzed for cytokine and chemokine concentrations by Bioplex assay.

^b^Analyte concentrations (pg/mL) are represented as the median value (n=8) per treatment group (two-independent experiments). The 95% confidence Interval (CI) is shown with the minimum and maximum value represented.

^c^The *p*-values represent comparisons generated from a Kruskal-Wallis with Dunn’s post-test comparing all groups. Bolded *p*-values are those that were found to be statistically significant, p < 0.05.

^d^‘LOD’ are those values that fall at or below the limit of detection.

**Table S2: Cytokine and chemokine concentrations in the brains of mice provided SKT20-LALA-PG^a^**

| **Analyte** | **Median (pg/mL) (95% CI)^b^** | | | ***p*-value^c^** | | |
| --- | --- | --- | --- | --- | --- | --- |
|  | Control | SKT20 | SKT20 LALA-PG | SKT20 v.  Control | SKT20 v. LALA-PG | LALA-PG v. Control |
| IL-6 | 512.3 (358.8 - 718.0) | 72.86 (14.5 - 419.6) | 395.3 (157.8 - 358.8) | **0.0005** | 0.09 | 0.36 |
| IFNγ | 370.8 (268.2 - 506.5) | 147.6 (71.3 - 332.8) | 386 (260.6 - 796.4) | **0.006** | **0.005** | >0.9999 |
| CXCL1 | 601.4 (490.5 -755.0) | 255.4 (147.5 - 354.0) | 478.8 (334.3-658.5) | 0.0002 | 0.02 | 0.66 |
| MCP-1 | 40,393 (32,409-55,394) | 7,490 (3,126 - 12,253) | 39,379 (16,455 - 48,525) | **0.0001** | **0.02** | 0.47 |
| MIP-1α/CCL3 | 3,299 (2,339-6,103) | 522.5 (244.1-807.7) | 2,068 (1302-3918) | **0.0001** | **0.04** | 0.54 |
| MIP-1β/CCL4 | 4,927 (3,916-7,338) | 1,121 (553.7-2,322) | 4,264 (3,243-6,662) | **0.0003** | **0.01** | 0.97 |
| RANTES/CCL5 | 13,340 (8,774-22,278) | 4,364 (3,086-6,583) | 9,186 (7,123-25,410) | **0.0002** | **0.01** | 0.77 |
| TNF-α | 82.2 (43.6-135.7) | 26.6 (20.4-45.2) | 54.9 (40.0-92.9) | **0.0009** | **0.02** | >0.9999 |
| IL-1β | 6.2 (5.3-7.4) | 2.6 (2.1-5.0) | 5.0 (4.1-7.6) | **0.0003** | **0.04** | 0.54 |
| IL-12p70 | 282.8 (147.5-1274) | 120.3 (66.0-302.5) | 260.1 (103.9-385.3) | **0.048** | 0.31 | >0.9999 |
| eotaxin | 840.5 (692.8-1023) | 375.5 (237.3-516.7) | 865.4 (715.7-1203) | **0.002** | **0.002** | >0.9999 |
| IL-12p40 | 2,659 (559.1-4,347) | 269.2 (114.4-1,668) | 994.1 (489.0-1,575) | 0.12 | 0.12 | 0.12 |
| G-CSF | 2,659 (559.1-4,347) | 269.2 (114.4-1,668) | 994.1 (489.0-1,575) | **0.0003** | 0.20 | 0.12 |
| GM-CSF | 70.5 (48.4-79.7) | 26.1 (19.8-55.7) | 56.1 (31.7-79.4) | **0.0022** | **0.0194** | >0.9999 |
| IL-1α | 36.6 (24.5-47.3) | 24.9 (20.1-40.8) | 36.4 (32.0-60.4) | 0.07 | **0.04** | >0.9999 |
| Il-2 | 17.5 (15.5-18.8) | 13.7 (11.3-15.6) | 16.4 (13.9-19.0) | **0.004** | **0.03** | >0.9999 |
| Il-3 | 15.0 (10.9-22.5) | 8.1 (5.4-16.6) | 14.7 (9.0-23.0) | **0.024** | **0.0441** | >0.9999 |
| IL-4 | 13.8 (9.7-20.4) | 7.4 (4.2-14.8) | 14.3 (8.5-18.6) | **0.005** | **0.03** | >0.9999 |
| IL-5 | 8.7 (4.1-11.3) | 4.1 (0.2-10.4) | 6.5 (1.1-8.6) | **0.04** | >0.9999 | 0.32 |
| IL-9 | 27.9 (20.4-45.6) | 18.2 (13.6-37.13) | 28.4 (17.9-43.6) | **0.046** | **0.048** | >0.9999 |
| IL-10 | 155.9 (118.3-191.5) | 70.22 (45.2-103.3) | 143.5 (118.7-171.9) | **0.0003** | **0.01** | >0.9999 |
| IL-13 | 138.4 (85.0-236.4) | 70.95 (32.0-172.3) | 142.3 (75.0-214.4) | **0.02** | 0.053 | >0.9999 |
| IL-17 | 7.5 (5.0-11.7) | 4.7 (3.4-9.7) | 7.5 (3.6-11.1) | 0.27 | 0.504 | >0.9999 |

^a^Mice were provided SKT20, SKT20-LALA-PG, or an isotype control one day prior to intranasal infection with TC-83. At 6 dpi, brains were collected, homogenized, and analyzed for cytokine and chemokine concentrations by Bioplex assay.

^b^Analyte concentrations (pg/mL) are represented as the median value (n=8) per treatment group (two-independent experiments). The 95% confidence Interval (CI) is shown with the minimum and maximum value represented.

^c^The *p*-values represent comparisons generated from a Kruskal-Wallis with Dunn’s post-test comparing all groups. Bolded *p*-values are those that were found to be statistically significant, p < 0.05.
